# Somatic copy-number alteration signatures reveal sex-divergent aging trajectories accelerated in cancer

**DOI:** 10.64898/2026.05.20.726597

**Authors:** Bingru Feng, Jun Xia, Yusi Fu

**Affiliations:** Center for Epigenetics and Disease Prevention, Institute of Biosciences and Technology, Texas A&M University, Houston, TX, 77030, USA; Center for Genomic and Precision Medicine, Institute of Biosciences and Technology, Texas A&M University, Houston, TX, 77030, USA

## Abstract

Somatic copy-number alterations (CNAs) accumulate with age and contribute to age-related pathologies, but their systematic characterization at single-cell resolution has been limited by the throughput-resolution trade-off in single-cell whole-genome sequencing.

Here, we developed ultra-CNA, a high-resolution single-cell analysis pipeline that extends CNA detection to 10-kb bin resolution and jointly profiles copy-number and single-nucleotide variation (SNV). Re-analyzing the Tasc-WGS dataset (Liu et al., 2022; previously analyzed at 200-kb resolution) of 32,526 lymphocytes from 16 healthy donors aged 0.7 to 79 years, we constructed a multi-dimensional CNA spectrum stratified by chromosomal context, copy-number state, size, and clonality. Small (<1 Mb), rare, predominantly loss-type CNAs accumulated progressively and stochastically with age. Sex-chromosome loss showed divergent kinetics: chromosome X loss cells in females accumulated at +0.10 percentage points per year, versus +0.03 for chromosome Y loss cells in males. Sex chromosome loss also had specific consequences for autosomal SNV burden: in younger donors, loss cells carried fewer autosomal SNVs than non-loss cells, whereas in older donors (>30 years), loss cells exceeded non-loss cells in both sexes. Female X-loss cells additionally exhibited elevated 45S rDNA copy number, supporting biologically distinct consequences of X loss and LOY. Clock-like SBS1 and SBS5 mutational signatures co-accumulated with age across both sexes.

Applying KL-divergence non-negative matrix factorization to the channelized CNA spectra, we constructed an aging clock validated by leave-one-sample-out cross-validation. Applied to a matched esophageal cohort, the clock detected accelerated aging from normal squamous epithelium through Barrett’s esophagus to esophageal adenocarcinoma, with cancer-associated spectra additionally enriched for large, highly clonal events. Ultra-CNA thus provides a scalable framework for quantifying somatic genomic aging from blood and for detecting accelerated aging in cancer.

## Introduction

Somatic mutations accumulate throughout life and generate genomic mosaicism within normal tissues, contributing to cellular dysfunction, clonal selection, tissue aging, neurodegeneration, and elevated cancer risk[1–4]. Among these, single-nucleotide variants (SNVs) have been most extensively characterized and shown to accumulate as a function of age across diverse human cell types[5–7]. Somatic copy-number alterations (CNAs) represent a complementary class of events that affect gene dosage across megabase-scale regions and arise through replication stress and chromosome mis-segregation[8–10]. Yet how CNAs accumulate during normal human aging, particularly at single-cell resolution and in coordination with point-mutation processes, remains far less understood. Prior single-cell studies in neurons reported anti-correlations between donor age and CNA frequency[11], but were limited by small cell populations and coarse detection resolution, precluding robust reconstruction of age-associated CNA trajectories.

A major bottleneck has been the throughput-resolution trade-off in single-cell whole-genome sequencing. Deep coverage protocols enable sensitive variant detection but scale poorly beyond hundreds of cells[12], while shallow high-throughput protocols capture thousands of cells but historically lacked the analytical resolution to detect small (<1 Mb), rare, non-clonal CNAs. Tasc-WGS, a Tn5-tagmentation-based single-cell whole-genome sequencing protocol, demonstrated the feasibility of processing ∼10,000 lymphocytes per day and was applied to a cohort of 16 cancer-free donors spanning 0.7 to 79 years of age, identifying large-scale (>2 Mb) CNAs and recurrent sex-chromosome aneuploidies at 200-kb bin resolution[13]. However, the original analysis focused on large events and did not integrate point-mutation profiles, leaving small, rare, low-clonality CNAs and their joint dynamics with SNV accumulation unexplored.

Sex chromosome loss is the most prominent age-associated CNA event in blood and merits particular attention in this context. In males, loss of chromosome Y (LOY) has been linked to aging, age-related disease, and elevated cancer risk[14–16]; in females, mosaic loss of chromosome X (mLOX) has been similarly associated with aging and with increased risk of myeloid and lymphoid leukemias[17,18]. Despite these associations, the cell-level prevalence of sex-chromosome loss, its relationship to autosomal CNA architecture, and its consequences for the broader somatic mutation landscape, including SNV burden and ribosomal DNA dosage, remain poorly characterized. Whether female X loss and male Y loss represent biologically equivalent processes or distinct phenomena with different downstream genomic consequences is also unresolved.

Molecular aging clocks have been developed from DNA methylation, transcriptomic, proteomic, and clinical-biomarker layers, and have substantially advanced the quantitative measurement of biological age[3,19]. Single-cell transcriptomic clocks add cellular-resolution age estimation but are sensitive to non-age-related sources of expression variation and typically remain tissue-specific[20–25]. DNA copy-number-based aging clocks, by contrast, remain largely unexplored, particularly in healthy human tissues and at single-cell resolution. Peripheral blood offers a minimally invasive and routinely accessible tissue source, and prior genome-wide association studies have used blood samples to characterize age-related mosaic chromosomal alterations and their associations with disease[26]. A CNA-based aging signature derived from healthy lymphocytes could therefore provide a scalable biomarker of molecular aging and a substrate for detecting accelerated aging in precancerous or malignant tissues.

Here, we developed ultra-CNA, a high-resolution single-cell analysis pipeline that extends CNA detection to 10-kb bin resolution and integrates copy-number and single-nucleotide variation analyses at single-cell scale. We applied ultra-CNA to our Tasc-WGS dataset comprising 32,526 lymphocytes from 16 healthy donors spanning 0.7 to 79 years[13], constructed a multi-dimensional CNA spectrum stratified by size, copy-number state, clonality, and chromosomal context, and identified progressive accumulation of small, rare, predominantly loss-type CNAs. Joint CNA-SNV analysis revealed sex-divergent dynamics: female X-loss cells accumulate faster than male LOY cells and show elevated ribosomal DNA copy number, and sex chromosome loss has sex-specific consequences for autosomal SNV burden, with clock-like SBS1 and SBS5 mutational signatures co-accumulating with age across both sexes (Fig. 1A). By applying KL-divergence non-negative matrix factorization to the channelized CNA spectra, we constructed an aging clock validated by leave-one-sample-out cross-validation, and applied it to a matched esophageal cohort spanning normal squamous epithelium, Barrett’s esophagus, and esophageal adenocarcinoma, demonstrating that the lymphocyte-derived signature detects accelerated aging in precancerous and malignant tissues (Fig. 1A).

**Figure 1.**
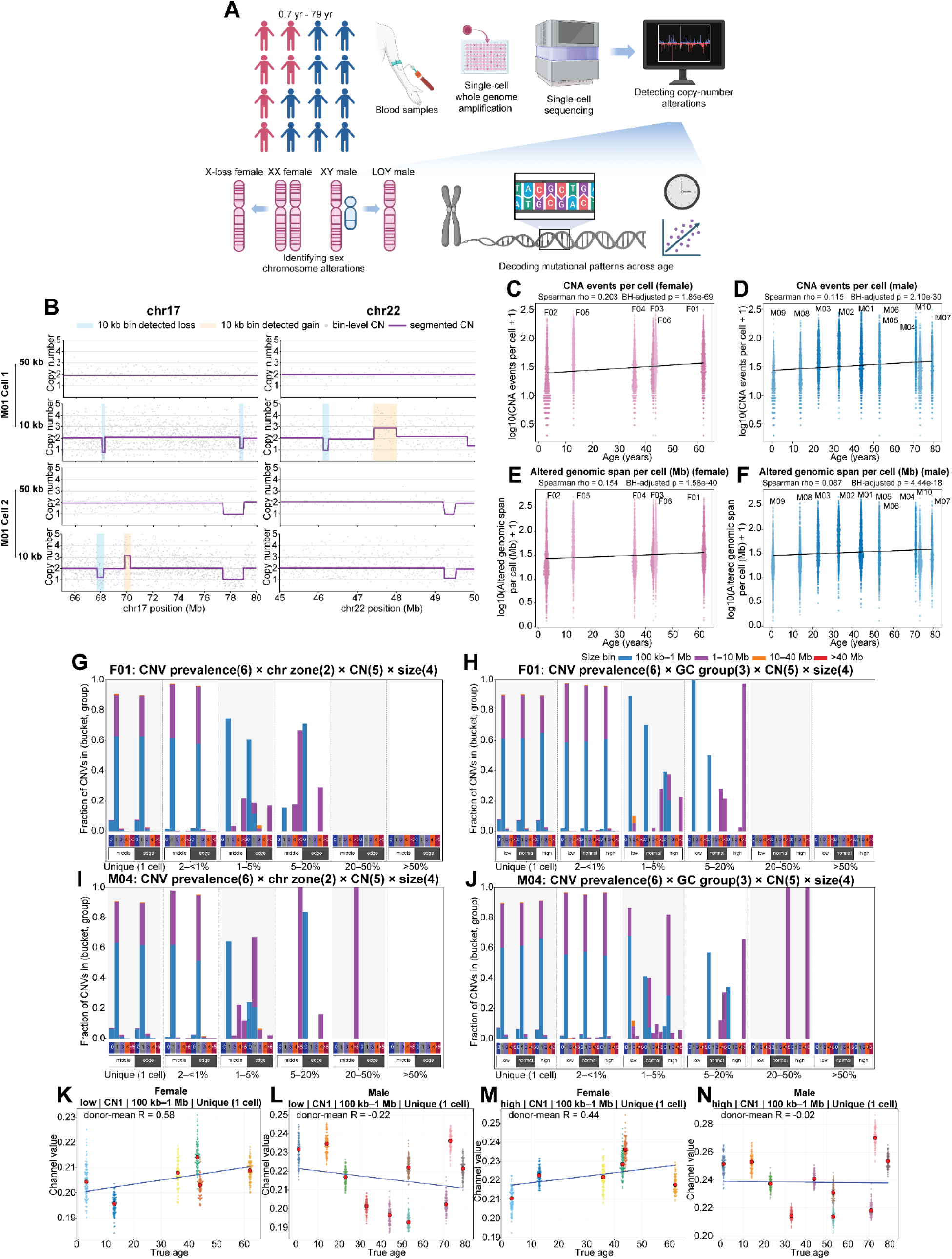
Ultra-CNA detects sex-divergent CNA accumulation in aging lymphocytes. (A) Overview of the ultra-CNA pipeline applied to single-cell whole-genome sequencing data from 16 donors spanning 0.7-79 years. (B) Representative single-cell CNA profile at 10-kb and 50-kb resolution. All plots show bin-level copy-number ratios (gray points) overlaid with the CBS-derived segmented signal (purple line). Grey lines mark integer copy-number levels. 10 kb bin detected copy-number loss and gain events are marked in blue and orange, respectively. (C-D) Number of somatic CNA events per cell versus donor age in females (C) and males (D). (E-F) Cumulative genomic span (Mb) affected by somatic CNAs per cell versus donor age in females (E) and males (F). (G-H) Representative CNA spectra for female donor F01: GC-content spectrum (G) and chromosomal arm-zone spectrum (H). Bars show the fraction of CNA events within each recurrence × GC (G) or recurrence × arm-zone (H) subgroup. Stacked colors denote event size (100 kb-1 Mb, 1-10 Mb, 10-40 Mb, >40 Mb). Tiles below the bars indicate copy-number state. (I-J) Same as (G-H), for representative male donor M04. (K-N) Sex-divergent CNA channels. Female (K, M) and male (L, N) channels (low GC, CN=1, 100 kb-1 Mb, singleton; K, L) and (high GC, CN=1, 100 kb-1 Mb, singleton; M, N) plotted against donor age. Each beeswarm shows bootstrap pseudosamples; red dots are donor medians; regression lines summarize the age trend. In C-F, each point is a single cell; pink = female, blue = male donors; black line, linear fit; Spearman’s rho and p-value shown. In G-J, only autosomal CNA events (chr1-22) are included. Full per-donor spectra in Figs. S2-S5.

## Results

### Small, rare, non-clonal CNAs accumulate progressively in aging lymphocytes

To characterize somatic CNA accumulation during aging at scale, we re-analyzed our single-cell whole-genome sequencing dataset generated by Tasc-WGS, originally processed at 200-kb bin resolution[13]. We developed ultra-CNA, a high-resolution computational pipeline that re-bins this dataset at 10-kb resolution and jointly profiles copy-number and single-nucleotide variation, with quality filtering, GC-bias correction, CBS segmentation, and integer copy-number estimation (Fig. 1B). Profiling at 10-kb resolution enabled detection of small CNA events that were not identifiable using 50-kb bins (Fig. 1B). The dataset comprised 32,526 single lymphocytes from 16 healthy donors (6 females, 10 males) spanning 0.7 to 79 years. After filtering for alignment rate >90%, autosomal MAD <0.507, and fraction of autosomal bins with CN≠2 <0.20, 17,503 cells were retained for downstream analysis. Both the frequency and burden of somatic CNAs increased with donor age (Figs. 1C-F, S1A-B), including the number of CNA events per cell, the cumulative genomic span affected, the fraction of the autosomal genome altered, and the number of chromosomes with CNAs per cell (Figs. 1C-F, S1C-F). Females showed steeper age-associated increases in both frequency and burden than males (Figs. 1C, E, S1C, E). Notably, when female and male donors were pooled into a single combined analysis (Figs. S1A-B), the age correlation and sex-specific differences were substantially attenuated compared to sex-stratified plots (Figs. 1C-F), underscoring that sex-specific trends are obscured when both sexes are analyzed jointly and motivating sex-separated analyses throughout this study.

To identify which classes of events drive this accumulation, we stratified CNAs by size, copy-number state, and clonality (Figs. S1G-I). The CNA landscape of healthy lymphocytes was dominated by small copy-number losses occurring predominantly as singletons (1 cell) or within rare subclones (2 cells - ≤1% of cells), with limited clonal expansion (Figs. S1G-I). Per-cell counts of the smallest (100 kb-1 Mb) size class increased with age (Figs. S1J-K), as did both loss and gain events (Figs. S1L-O) and singleton events specifically (Figs. S1P-Q). The dominance of singletons implies that most CNAs arise de novo in individual cells rather than through clonal expansion, consistent with stochastic mutational processes rather than positive selection in healthy lymphocytes, which parallels reported age-associated increases in private somatic mutations genome-wide[27]. As with global CNA accumulation, these subclasses showed steeper positive age trends in females.

### Sex-divergent CNA spectra define distinct aging architectures

To systematically resolve sex-specific differences in CNA architecture, we developed a multi-dimensional CNA spectrum framework. Motivated by analogous SBS-signature strategies, in which sparse high-dimensional mutation matrices are compressed into interpretable lower-dimensional components[28], we constructed two complementary spectrum representations per donor: a 240-dimensional chromosomal arm-zone spectrum stratified by arm position (edge, middle), copy-number state (CN = 0, 1, 3, 4, >5), event size (100 kb-1 Mb, 1-10 Mb, 10-40 Mb, >40 Mb), and clonality (singleton, 2 cells-<1%, 1-5%, 5-20%, 20-50%, >50%) (Figs 1G,I, S2A-E, S3A-I); and a 360-dimensional GC-content spectrum stratified by local GC category (low, normal, high) in combination with copy-number state, size, and clonality (Figs 1H,J, S4A-E, S5A-I). CNA events within each donor were clustered using a graph-based maximal-clique approach requiring ≥90% reciprocal overlap between segments of the same integer copy-number state on the same chromosome, then mapped to their corresponding channels and normalized within clone-frequency bins to produce a compositionally stable feature vector.

Representative spectra illustrated the sex-specific architecture. We selected F01 (62 years, eldest female) and M04 (71 years, closest-aged elderly male) as representative donors for direct visual comparison of CNA profiles between sexes at advanced age. Female donor F01 showed a spectrum dominated by small, low-clonality loss events in both arm-zone and GC-content representations (Figs. 1G-H), while male donor M04 displayed a qualitatively similar architecture with slightly greater clonal expansion in 1-10 Mb events (Figs. 1I-J). Direct comparison of age-associated channels between sexes revealed sex-divergent patterns: several classes, including (low GC, CN=1, 100 kb-1 Mb, singleton) and (high GC, CN=1, 100 kb-1 Mb, singleton), correlated positively with age in females, while in males the corresponding channels showed either negative correlation or little to no correlation (Figs. 1K-N). These divergent channels indicate that female and male lymphocytes differ not only in the overall rate of CNA accumulation but also in the qualitative remodeling of the CNA landscape with age. Together, these data establish that aging in healthy lymphocytes is associated with progressive accumulation of rare, small, predominantly loss-type CNAs, with sex-divergent trajectories across specific subclasses, and underscore the need for high-resolution single-cell tools such as ultra-CNA to capture the low-frequency, small-scale events that define this landscape and to support sex-specific clock construction downstream.

### Sex chromosome loss is a major axis of age-associated CNA accumulation

To systematically characterize genome-wide CNA patterns at the sample level, we compiled per-donor single-cell copy-number profiles into hierarchically clustered heatmaps (Figs. 2A-D, S6A-D, S7A-H). Copy-number losses predominated over gains across the cohort (Figs. 2A-D). Although most age-associated CNAs were small and autosomal, single-cell heatmaps revealed a distinct class of large-scale events affecting sex chromosomes in older donors: a subset of cells in aged female donors exhibited copy-number loss spanning the entire X chromosome, and a corresponding subset in aged male donors showed Y chromosome loss (Fig. 2B, D). Among all large-scale CNA events detected, sex chromosome loss showed the strongest and most consistent association with donor age.

**Figure 2.**
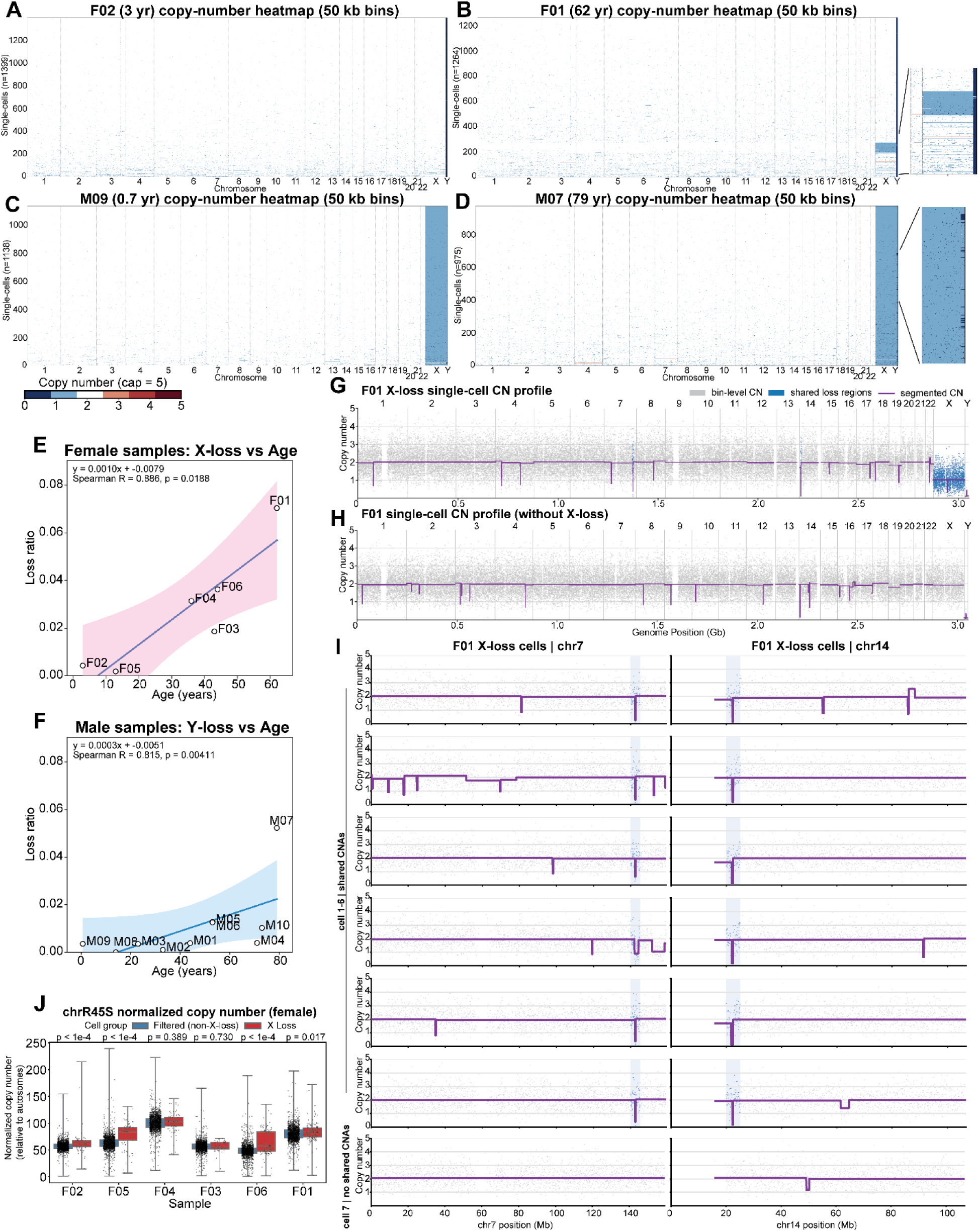
Sex chromosome loss and shared autosomal CNAs in X-loss cells. (A-D) Single-cell copy-number heatmaps for female donors F02 (A), F01 (B) and male donors M09 (C), M07 (D) shown at 50-kb display bins (color scale capped at 5). Rows, single cells; columns, genome-ordered bins. Cells hierarchically clustered. Insets show sex-chromosome zoom-ins for X-loss (B) and LOY (D) cells. (E-F) Fraction of LOY cells in males (E) and X-loss cells in females (F) versus donor age. Each point is one sample; line, linear fit; shading, 95% CI; regression equation, Spearman’s r, and p-value shown. (G-H) Whole-genome copy-number profiles of an F01 X-loss cell (G) and a non-X-loss cell (H). (I) Per-chromosome copy-number profiles of six representative F01 cells showing recurrent losses on chromosomes 7 and 14. Blue points, recurrent loss regions; purple line, segmented profile. (J) Normalized chrR45S copy number in female X-loss versus read-depth-matched non-X-loss cells, per donor. Boxes, median and IQR; whiskers, range; points, single cells. P-values per donor.

To quantify sex chromosome loss, we defined X-loss cells in females as those in which >80% of chromosome X bins had copy-number values <2, and LOY cells in males as those in which >80% of chromosome Y bins had copy-number values <1. In females, the fraction of X-loss cells increased significantly with age at a steeper rate (+0.10 percentage points per year; Fig. 2E). The fraction of LOY cells also increased with age in males (slope: +0.03 percentage points per year; Fig. 2F), consistent with LOY as a well-established form of age-related clonal hematopoietic mosaicism[14,16]. This suggests more rapid age-associated accumulation of X loss than LOY in this cohort.

Representative single-cell copy-number profiles from F01 confirmed the X-loss phenotype (Fig. 2G). An F01 X-loss cell showed broad chromosome X copy-number loss (CN=1), whereas an F01 cell without X loss retained chromosome X copy number (CN=2) (Figs. 2G-H). We further examined autosomal CNA profiles across F01 X-loss cells to identify shared copy-number events indicating common clonal origin among sex-chromosome-loss cells. Per-chromosome views of six representative F01 cells (cells 1-6) revealed losses on chromosomes 7 and 14, supporting clonally expanded autosomal CNA events in a subset of F01 X-loss cells (Fig. 2I). To exclude the possibility that shared autosomal CNAs could arise by chance in non-loss cells, we compared X-loss or LOY cells with donor-matched non-loss cells selected for similar total read counts; the matched non-loss set contained the same number of cells as the corresponding X-loss or LOY group. We performed the analysis in donors with >10 sex-chromosome-loss cells (F04, F03, F06, F01, M10, M07). Among these donors, sex-chromosome-loss cells showed more shared autosomal loss and gain bins compared to read-depth-matched non-loss cells (Figs. S8A-L). We further quantified the total number of loss-cell-enriched and non-loss-cell-enriched bins across donors and across five random matched non-loss comparisons per donor, and found that sex-chromosome-loss cells had significantly more shared autosomal loss-CNA and gain-CNA bins than matched non-loss cells, using recurrence-fraction cutoffs of 0.15 for loss CNAs and 0.05 for gain CNAs (Figs. S8M-N). The labeled regions indicate shared autosomal CNA events among X-loss or LOY cells (Figs. S8A-L), supporting shared clonal expansion among sex-chromosome-loss cells rather than random, cell-specific copy-number changes. Together, these findings identify sex chromosome loss as a major axis of age-associated large-scale CNA accumulation in lymphocytes, with distinct kinetics between sexes.

### Female X-loss cells exhibit elevated 45S ribosomal DNA copy number

To investigate whether sex chromosome loss is associated with broader copy-number remodeling beyond the standard chromosomes, we estimated relative 45S rDNA abundance from read depth mapped to the 45S rDNA contig (chrR45S), normalized to autosomal read depth per cell. In female donors, X-loss cells showed higher normalized chrR45S signal than non-X-loss cells from the same donor, with this difference reaching statistical significance in four of six female donors (Fig. 2J). This pattern was not observed between LOY and non-LOY cells in male donors (Fig. S9A). These results suggest that X chromosome loss in female lymphocytes is associated with a distinct genomic state including elevated rDNA copy number, potentially reflecting compensatory genomic remodeling. The absence of a corresponding rDNA signal change in male LOY cells further supports the conclusion that X loss and LOY represent biologically distinct processes with different downstream genomic consequences.

### Somatic SNV burden scales with chromosome length and is reduced on sex chromosomes

Having identified progressive accumulation of rare and small structural variants in our cohort, we next examined single-nucleotide variations (SNVs) to assess whether point-mutation patterns also reflected aging-associated processes. To characterize the genome-wide landscape of somatic point mutations at single-cell resolution, we first examined the relationship between covered chromosome length and mean per-cell homozygous SNV burden in each donor. In both females and males, mean per-cell homozygous SNV burden was strongly positively correlated with covered chromosome length (Pearson r = 0.97, females, Fig. 3A; r = 0.96, males, Fig. 3B), indicating that longer chromosomes accumulate proportionally more somatic mutations, consistent with a stochastic mutational process across the genome.

**Figure 3.**
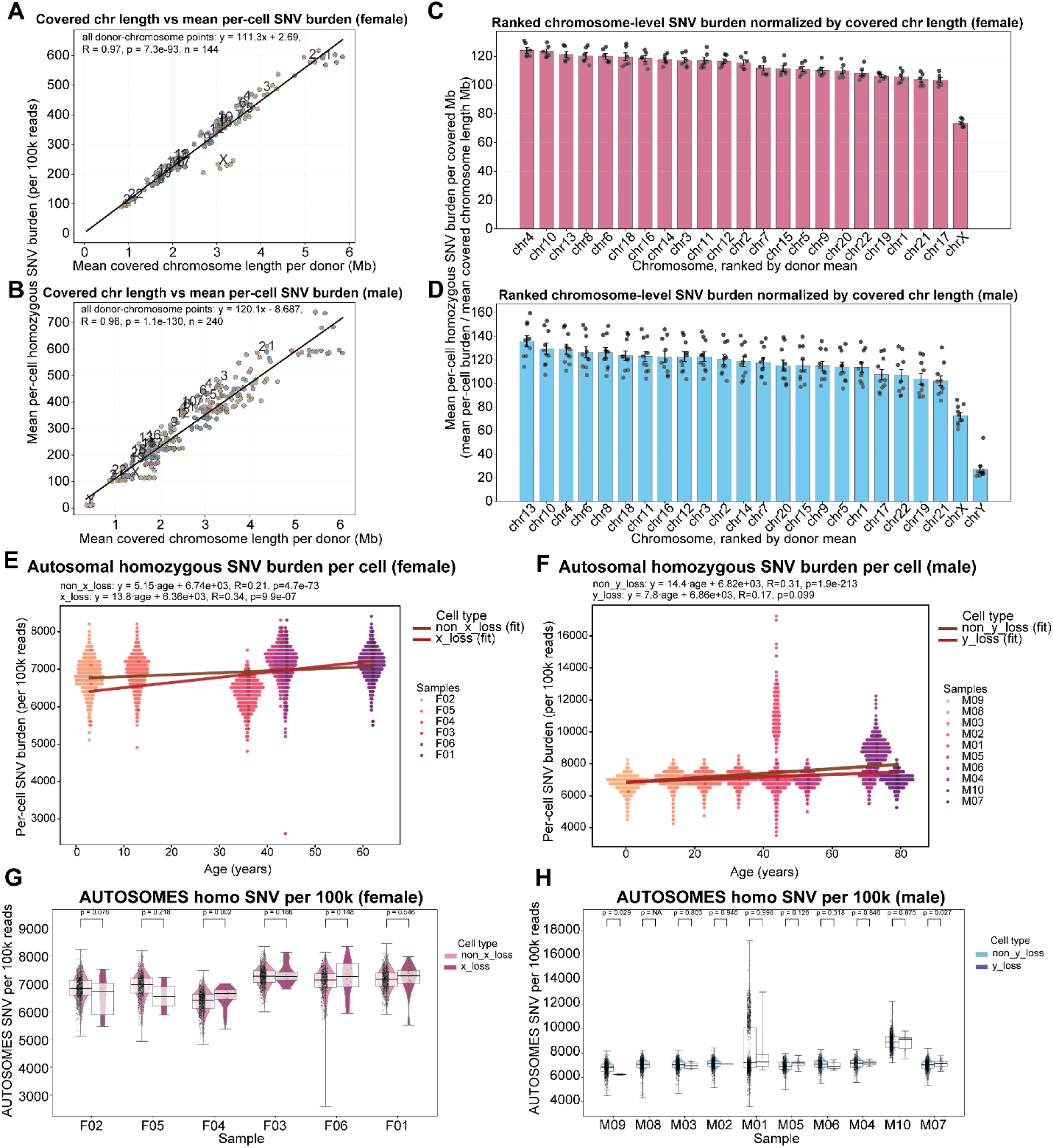
Genome-wide somatic SNV burden in females and males. (A-B) Mean per-cell homozygous SNV burden versus covered chromosome length in females (A) and males (B). Each point is one donor-chromosome pair; black line, linear fit; Pearson r shown. (C-D) Chromosomes ranked by mean homozygous SNV burden per covered Mb in females (C) and males (D). Bars, mean across donors; error bars, SEM; points, individual donors. (E-F) Per-cell autosomal homozygous SNV burden versus donor age in females (E) and males (F). Each dot, one cell; lines, linear fit. (G-H) Distribution of per-cell autosomal homozygous SNV burden in non-loss versus loss cells, per donor (G: female non-X-loss vs. X-loss; H: male non-LOY vs. LOY). Violins, full distribution; embedded boxes, median and IQR. Per-donor comparison shown above each pair. SNV burden normalized per 100,000 reads, adjusted by alignment rate.

To identify chromosomes deviating from this expectation, we normalized mean homozygous SNV burden by covered chromosome length and ranked chromosomes accordingly (Figs. 3C-D). In females, chromosome X exhibited a notably lower normalized SNV burden than autosomes (Fig. 3C). In males, both chromosomes X and Y ranked among the lowest in normalized SNV burden (Fig. 3D). Because male chrX and chrY are hemizygous, we applied a ploidy-correction factor of 2 to rescale their normalized burden to a diploid-equivalent scale; even after this adjustment, chromosome Y remained notably depleted in SNV burden relative to autosomes, suggesting that factors beyond simple hemizygosity-related detection loss may shape the reduced SNV burden on sex chromosomes, including selective constraint or preferential elimination of cells with excess sex-chromosome mutations in both sexes.

### Autosomal SNV accumulation in sex-chromosome-loss cells diverges with age

Having established genome-wide SNV patterns, we next asked whether cells with sex chromosome loss exhibit altered somatic mutation dynamics. In both female and male donors, autosomal homozygous SNV burden increased with age regardless of sex-chromosome-loss status (Figs. 3E-F), confirming that age-associated SNV accumulation is a universal feature of aging lymphocytes.

However, the relationship between sex chromosome loss and point-mutation burden was not static across the lifespan. Per-donor comparisons between sex-chromosome-loss and non-loss cells revealed an age-dependent transition (Figs. 3G-H): in younger donors (<30 years), sex-chromosome-loss cells exhibited lower autosomal SNV burden than non-loss cells from the same individual, whereas in older donors (>30 years), this relationship reversed, with loss cells carrying higher autosomal SNV burden than non-loss cells from the same donor. This transition was observed in both female X-loss versus non-X-loss cells (Fig. 3G) and male LOY versus non-LOY cells (Fig. 3H). Because lymphocytes inherit most of their SNV burden from the hematopoietic stem cell of origin, the lower burden of loss cells in young donors likely reflects a selective bottleneck at the hematopoietic stem cell level: only stem cells with low pre-existing mutation burden tolerate sex chromosome loss and successfully expand. These loss-bearing stem cell clones subsequently accumulate mutations at an accelerated rate, eventually surpassing non-loss cells, consistent with our previous findings in aged male cancer patients in which LOY cells showed broader genome instability than non-LOY cells[29].

Comparing SNV accumulation rates between sex-chromosome-loss populations across sexes, female X-loss cells showed a higher autosomal SNV accumulation rate (13.8 SNVs per 100,000 reads per year) than male LOY cells (7.8 SNVs per 100,000 reads per year) (Figs. 3E-F). This difference may reflect the greater functional consequence of X-chromosome loss in females: the X chromosome harbors numerous genes involved in DNA repair, chromatin maintenance, and genome stability (e.g., ATRX, HDAC6, SMC1A)[30–32], and loss of one X copy may disrupt dosage of these regulators, triggering broader genomic instability. By contrast, loss of the gene-poor Y chromosome may impose fewer immediate consequences on genome maintenance, resulting in a comparatively modest SNV accumulation trajectory in LOY cells.

Notably, chromosomes with the highest normalized SNV burden (chr13, chr4, and chr10; Figs. 3C-D) exhibited age-associated accumulation trends closely mirroring those for total autosomal burden (Figs. S9B-G), indicating that the divergent mutation dynamics in chromosome-loss cells are not driven by a single outlier chromosome but reflect a genome-wide phenomenon.

In male LOY cells, a distinct pattern emerged on chromosome X: while non-LOY cells showed the expected age-associated increase in chrX homozygous SNV burden, LOY cells displayed a non-significant downward trend with age (Fig. S9H). This is notable given our previous observations in cancer, where LOY was associated with compensatory chromosome X copy-number gain[29]. In healthy lymphocytes, however, neither X copy-number gain nor increased chrX SNV burden was detected, suggesting an alternative strategy: LOY cells may face selective pressure to preserve chromosome X integrity, given that chrX harbors numerous genes homologous to Y-linked counterparts (e.g., UTX/UTY, DDX3X/DDX3Y, ZFX/ZFY)[33]. In the absence of Y-linked paralogs, maintaining a low-mutation X chromosome may become critical for cell survival, potentially through enhanced repair of X-linked sequences. This would explain both the reduced chrX SNV signal in LOY cells (Fig. S9H) and the observation that surviving LOY cells nonetheless accumulate autosomal mutations in older age (Fig. 3H): genomic maintenance may be preferentially directed toward chromosome X at the expense of autosomal fidelity.

Together, these findings reveal that sex chromosome loss has sex-specific and chromosome-specific consequences for the broader somatic mutation landscape, while the general age-associated increase in autosomal SNV burden is preserved across all cell populations regardless of sex chromosome status.

### Clock-like SBS1 and SBS5 mutational signatures accumulate with age

Refitting of donor-level single-base substitution (SBS) 96 mutational catalogs against COSMIC reference signatures identified contributions from established clock-like signatures[28,34], including SBS1 and SBS5. Observed and reconstructed SBS96 profiles for representative eldest donors, F01 (62 years, female) and M07 (79 years, male), confirmed reliable reconstruction of both SBS1 and SBS5 components (Figs. 4A-C, S10A-C), and cohort-level signature attribution across all female and male donors verified that age-associated mutational processes are represented throughout this aging cohort (Figs. S10D-G). SBS1 arises from spontaneous or enzymatic deamination of 5-methylcytosine at CpG dinucleotides and accumulates in a cell-division-dependent manner[35], whereas SBS5 is a broadly distributed clock-like signature of unknown etiology that increases with age across many tissue types[36] (Figs. 4D-E). SBS46 and SBS54 were excluded from subsequent analysis as possible sequencing artifacts[28] (Figs. 4D-E). Three male donors (M08, M01, and M10) showed additional signature contributions beyond the SBS1/SBS5 pattern (Figs. 4D-E). SBS3 is associated with defective homologous recombination-mediated DNA damage repair, typically linked to BRCA1/BRCA2 mutations, and produces a characteristic flat distribution across the 96 trinucleotide contexts[28]. SBS37 and SBS102 are rare signatures of unknown etiology that have been observed in a limited number of samples[28]. The presence of these additional signatures in a subset of male donors suggests inter-individual variation in active mutational processes beyond the universal clock-like SBS1 and SBS5, though their biological significance in healthy lymphocytes remains unclear.

**Figure 4.**
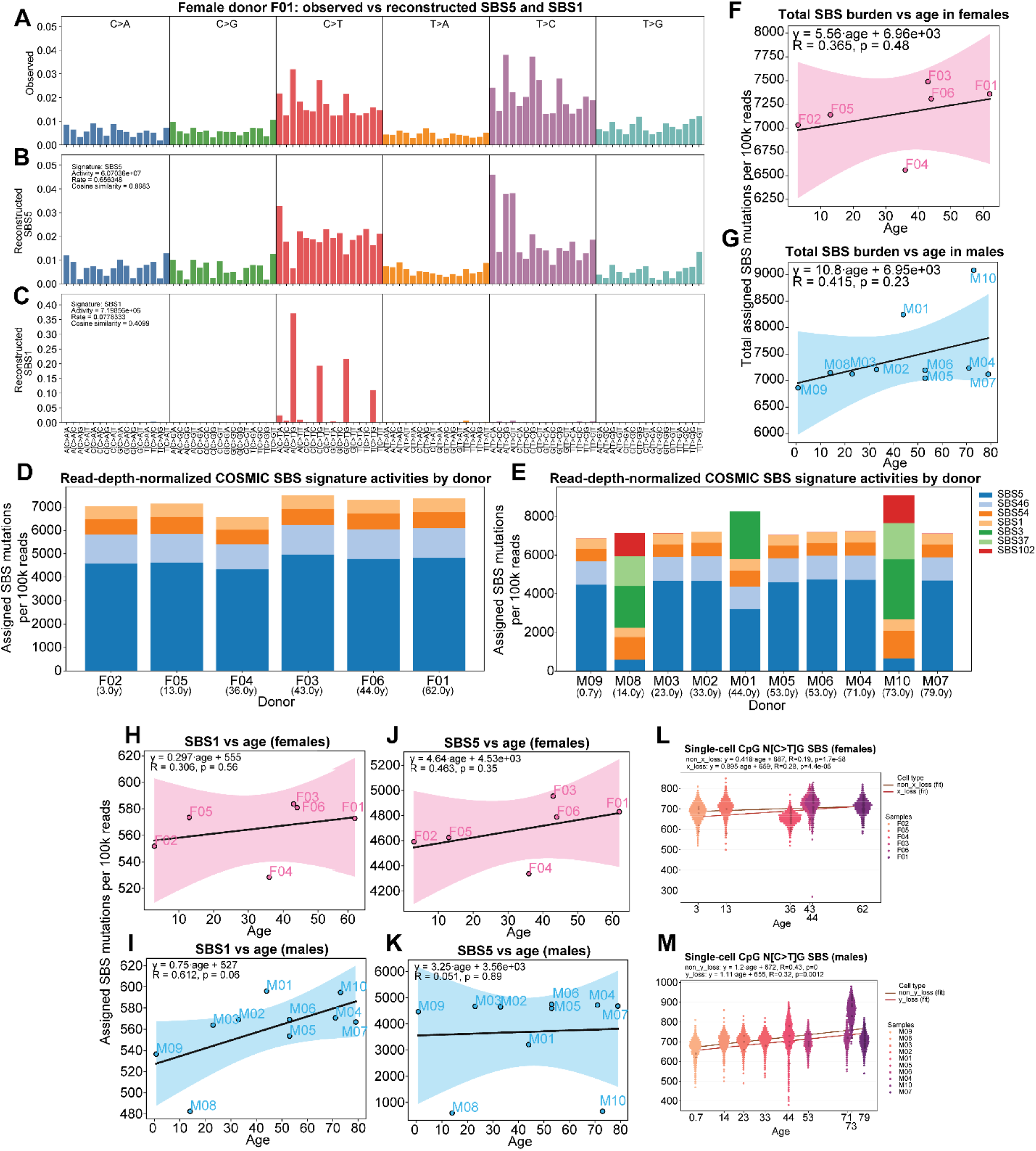
SBS96 mutational spectra and top enriched signatures. (A) Observed pooled SBS96 spectrum in F01. (B) Reconstructed SBS96 spectrum from fitted SBS5 contributions in F01. (C) Reconstructed SBS96 spectrum from fitted SBS1 contributions in F01. (D-E) Normalized COSMIC SBS signature activities per donor in females (D) and males (E), ordered by age. (F-G) Normalized total SBS burden versus age in females (F) and males (G). (H-K) Age associations of top SBS signatures: SBS1 (H) and SBS5 (J) in females; SBS1 (I) and SBS5 (K) in males. Lines, sex-specific linear regression; Pearson r and p-value shown. (L-M) Single-cell CpG (N[C>T]G) SBS burden versus age (L: females, M: males). Each dot represents one cell; lines, linear fit. Activities normalized by sequencing depth and alignment rate.

Total SBS burden and SBS1 burden, normalized by each donor’s read depth, both showed positive age trends in females and males, with higher accumulation in males (Figs. 4F-I). For the dominant signature, SBS5, activities were generally higher and accumulated more aggressively in females than in males (Figs. 4J-K), suggesting sex differences in SNV accumulation dynamics. To evaluate single-cell-level SBS1 activity, we quantified spontaneous deamination of 5-methylcytosine at CpG dinucleotides per cell by counting homozygous SNVs in CpG C>T SBS96 contexts (N[C>T]G), normalized by read depth and alignment rate. Normalized single-cell SBS1 activity increased with age in both sexes, supporting progressive accumulation of age-related CpG deamination at single-cell resolution (Figs. 4L-M). The strongest age-associated increase was observed in male non-LOY cells, followed by male LOY cells, female X-loss cells, and female non-X-loss cells (Figs. 4L-M), indicating that age-related SBS1 accumulation varies by sex and sex-chromosome-loss state and that sex chromosome mosaicism is associated with distinct mutational dynamics.

### Stratification by size and clonality reveals clock-like CNA accumulation

Having established the CNA architecture of aging lymphocytes, we asked whether the same architecture appears in other healthy tissues and how it compares with cancer. We applied the CNA spectrum framework to two additional cohorts: an external single-cell neuron dataset[11] (Figs. 5A-B); and a matched esophageal cancer cohort comprising normal squamous epithelium (N), Barrett’s esophagus (BE), and esophageal adenocarcinoma (EAC) specimens[29] (Figs. 5C-D, S11A-F, S12A-F), and compared them against the healthy lymphocyte cohort[13] (Figs. 1G-J, S2A-E, S3A-I, S4A-E, S5A-I).

**Figure 5.**
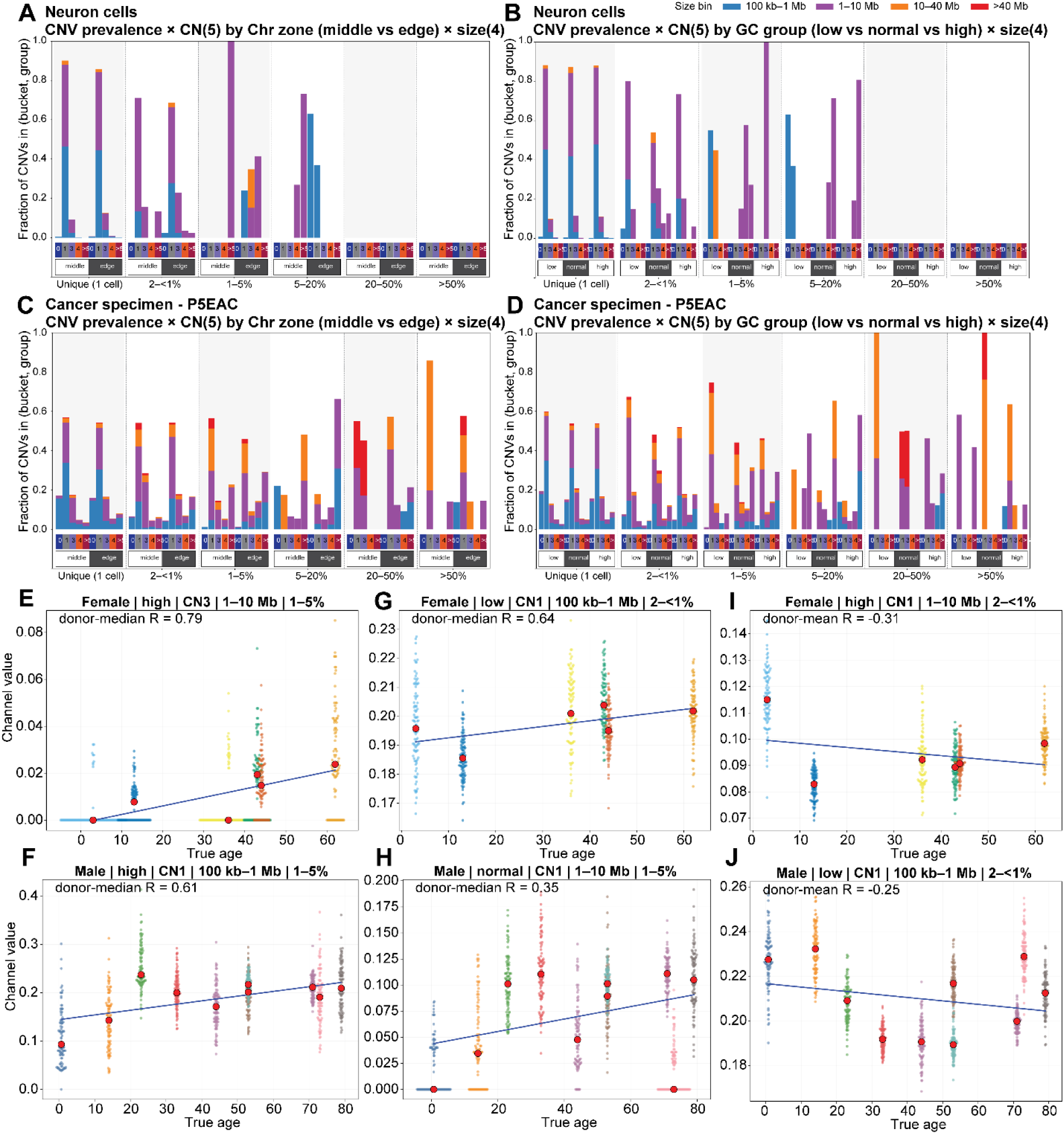
CNA spectra across cohorts and age-associated channels. (A-B) GC-content (A) and arm-zone (B) spectra of a representative external male neuron donor. Layout as in Figure 1G, H. (C-D) GC-content (C) and arm-zone (D) spectra of a representative EAC sample (P5EAC). Layout as in Figure 1G, H. (E-H) Top positively weighted CNA channels in the age-associated NMF component. Each panel shows one high-weighted channel plotted against donor age. Beeswarms, bootstrap pseudosamples; red dots, donor medians; regression lines summarize the age trend. (I-J) Top negatively weighted channels from the component with the strongest negative age correlation. Layout as in E-H. Only autosomal CNA events (chr1-22) are included. Full per-donor spectra in Figs. S11-S12.

Healthy neurons recapitulated the architecture of healthy lymphocytes: CNA events were predominantly small (100 kb-1 Mb), low copy-number losses (CN = 1), largely confined to the lowest clone-frequency bins (singleton and 2-<1%), indicating that they arise randomly in individual cells with limited clonal expansion (Figs. 1G-J, 5A-B). This conservation across two unrelated tissues, one post-mitotic and one mitotic, indicates that the small, rare, non-clonal CNA architecture is a general feature of healthy human aging rather than a lymphocyte-specific phenomenon.

In sharp contrast, EAC samples were enriched for large (>10 Mb), highly recurrent CNA events spanning multiple clone-frequency bins, reflecting the selective clonal expansions characteristic of cancer progression (Figs. 5C-D, S11A-F, S12A-F). The CNA spectrum thus discriminates two qualitatively distinct biological strategies: continuous accrual of small, stochastic, non-clonal events during normal aging versus large, clonal events that dominate oncogenic evolution.

To identify the CNA subclasses that most strongly track chronological age across donors, we examined the top age-correlated channels in the healthy cohort. Small CNAs, particularly those below 1 Mb and in the 1-10 Mb range, showed the strongest age-associated increase, occurring predominantly in rare subclones (clone frequency <5%) and indicating continuous accumulation throughout the lifespan rather than large clonal expansion (Figs. 5E-H). We also identified top anti-correlated channels, including 1-10 Mb rare subclones with CN = 1 in females and 100 kb-1 Mb rare subclones with CN = 1 in males (Figs. 5I-J). These negatively age-associated events were also predominantly small and low-clonality. Together, the bidirectional age trajectories of positively and negatively associated CNA classes support a clock-like model of CNA accumulation in which specific subclasses change in a continuous, age-dependent manner during normal aging, and provide the candidate feature space from which a quantitative aging clock can be learned.

### NMF identifies a CNA aging clock correlated with chronological age

While individual CNA channels showed age associations, these relationships were correlated and high-dimensional, motivating an unsupervised decomposition approach to identify compact latent CNA programs that best capture the aging trajectory. We applied KL-divergence non-negative matrix factorization (KL-NMF) to the pseudosample-by-channel feature matrices across candidate component numbers (K = 2, 3, 4, 5, and 6), using multiplicative-update optimization under Kullback-Leibler divergence loss. This yielded, for each K, a component-by-channel signature matrix and a matrix of pseudosample exposure fractions. Donor-level median exposures were calculated by averaging across 100 bootstrap pseudosamples (500 single cells each) per donor, separating biological signal from cell-sampling noise.

To isolate the component most informative of chronological age, we evaluated each component across all tested K values by correlating donor-median exposure with chronological age and ranking components by strength of positive age association (Figs. S13A). The final model was selected as the K achieving the highest donor-median R² on the age calibration regression, with the top-ranked age-positive component designated the CNA aging clock component (Figs. S13A). The trained clock comprises (i) the NMF component signatures (Figs. S13A), (ii) the identity of the age-associated clock component (Figs. 6A, S13B-E), and (iii) the linear calibration coefficients (slope and intercept) mapping that component’s donor-median exposure to chronological age (Figs. 6B). Leave-one-sample-out (LOSO) cross-validation in the female cohort confirmed robust age prediction in both sexes (Figs. S12F-H).

**Figure 6.**
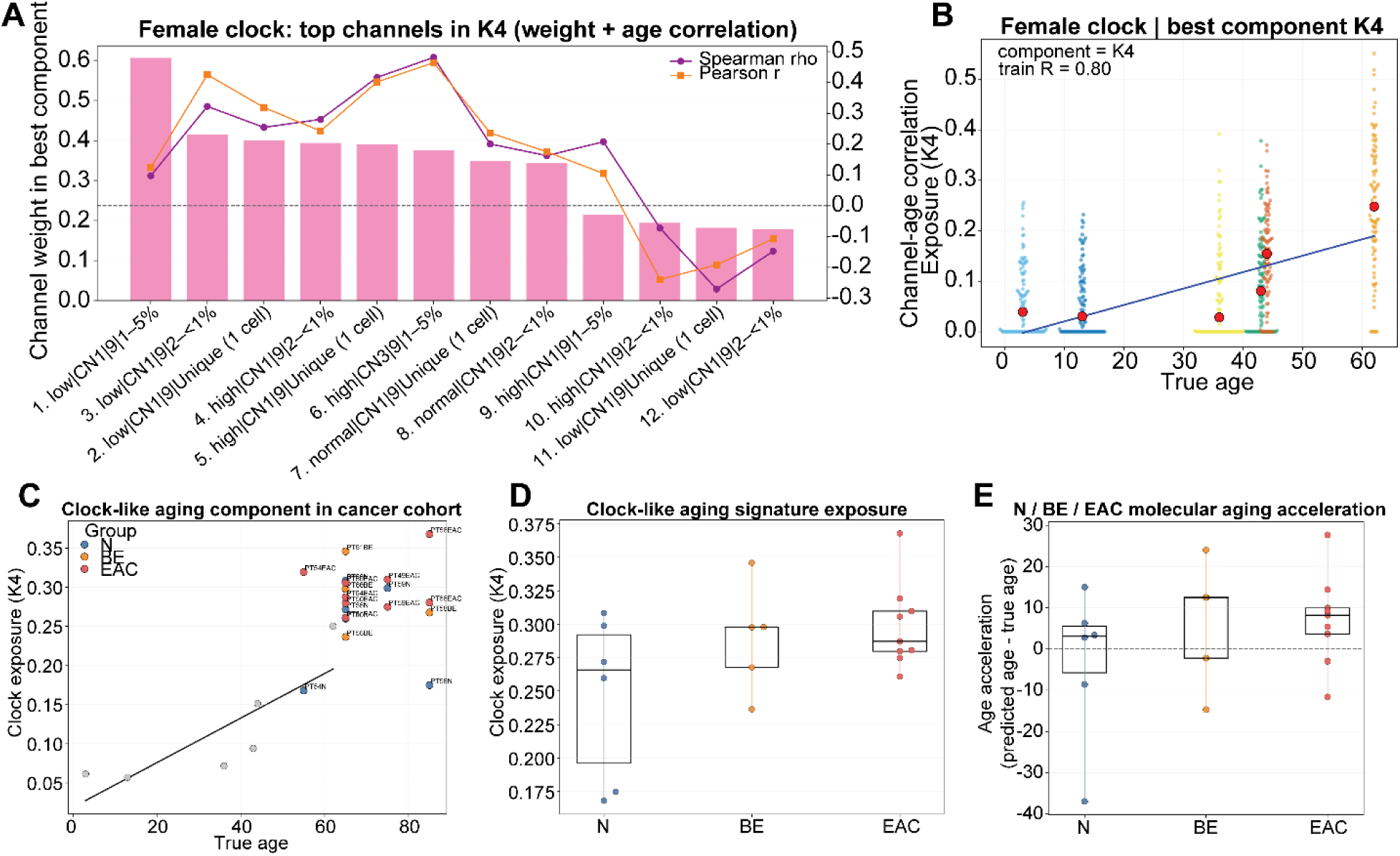
Training-cohort performance and external validation of the CNA aging clock. (A) Top-weighted CNA channels in the selected aging-clock component for female cohort. Bars, channel weights; overlaid lines, Spearman’s rho (purple) and Pearson’s r (orange) versus age. Channel labels: GC bin, CN state, size class, clone-frequency bucket. (B) Clock-component exposure versus chronological age in the female training cohort. Beeswarms, bootstrap pseudosamples; central markers, donor medians; line, linear fit; Pearson r and R² shown. (C) Clock-component exposure versus chronological age in the cancer cohort. Gray points, training donors; black line, training-cohort regression. Colored points, external esophageal samples (green, N; blue, BE; red, EAC). Upward displacement indicates elevated aging-associated CNA signature. (D) Distribution of clock-component exposure by histologic group (N, BE, EAC). Boxes, median and IQR; whiskers, range; points, individual samples. (E) Age acceleration (predicted minus chronological age) by histologic group. Layout as in (D); dashed line, no acceleration.

Because healthy lymphocytes exhibit sex-specific CNA patterns, including distinct sex-chromosome-loss and SNV dynamics, we implemented a sex-specific aging clock. This strategy ensures that male-associated CNA features do not confound the female age signal and produces a clock calibrated to the biological aging trajectory of females. The female clock demonstrated strong concordance between NMF component exposure and chronological age within the training cohort (Fig. 6B), establishing that CNA spectral decomposition can recover a quantitative, clock-like molecular readout of aging from single-cell copy-number profiles.

### The CNA aging clock detects accelerated aging from normal squamous epithelium through Barrett’s esophagus to esophageal adenocarcinoma

To test whether the lymphocyte-derived CNA aging clock detects altered molecular aging in disease contexts, we applied the trained clock to an independent cohort of matched esophageal specimens comprising normal squamous epithelium (N), Barrett’s esophagus (BE), and esophageal adenocarcinoma (EAC). Each external specimen was processed through the identical channelization pipeline used in training: CNA events were clustered per sample using the same graph-based maximal-clique approach with ≥90% reciprocal overlap, annotated by GC-content bin, copy-number category, event size, and clone-frequency bucket, and aggregated into the identical fixed-length channel feature vector. Each external sample vector was then projected onto the trained NMF component signatures using non-negative least-squares to obtain exposure fractions, and predicted ages were computed from the training-cohort linear calibration applied to the selected clock component’s exposure.

Projection of external esophageal specimens onto the training-cohort aging trajectory revealed a progressive increase in clock-component exposure from normal tissue through Barrett’s esophagus to esophageal adenocarcinoma (Figs. 6C, D). Most normal squamous epithelial specimens clustered near the expected aging trajectory defined by age-matched healthy lymphocyte donors, indicating exposures consistent with their chronological age. In contrast, BE and EAC specimens showed elevated clock-component exposure relative to age-matched normal tissue (Fig. 6D). This was reflected in a systematic molecular age acceleration from N to BE or EAC, defined as predicted age minus chronological age (Fig. 6E), indicating that premalignant and malignant esophageal lesions carry a CNA burden resembling that of a considerably older individual.

As described above, BE and EAC spectra were architecturally distinct from healthy aging tissues (Figs. 5C-D, S11A-F, S12A-F). The CNA aging clock, trained on healthy lymphocytes, is therefore not simply detecting the same aging process at an accelerated rate in cancer; rather, it is detecting an overall elevation in the genomic instability signature that partially overlaps with the aging-associated CNA program but is driven by fundamentally different biology.

Together, these results demonstrate that the CNA aging clock, derived entirely from healthy peripheral blood lymphocytes, can be applied to independent epithelial tissue specimens and detects a meaningful gradient of molecular aging acceleration across the histologic spectrum from normal esophageal mucosa to invasive adenocarcinoma. These findings suggest that CNA-based aging signatures may provide a useful framework for quantifying molecular aging acceleration in premalignant lesions and for stratifying disease risk in clinical specimens.

## Discussion

In this study, we developed ultra-CNA and used it to systematically characterize somatic CNA accumulation across the human lifespan at single-cell resolution. Aging in healthy lymphocytes is dominated by small, rare, predominantly loss-type CNAs accumulating in a non-clonal manner, with sex-divergent trajectories across specific CNA subclasses. Sex-chromosome loss is itself sex-divergent: female X-loss cells accumulate 3-fold faster than male LOY cells, carry elevated 45S rDNA copy number, and display distinct autosomal SNV burden dynamics relative to non-loss cells, with loss cells in both sexes exceeding non-loss cells after ∼30 years of age. KL-NMF decomposition of the CNA spectrum yields a female aging clock that detects accelerated molecular aging from normal squamous mucosa through Barrett’s esophagus and esophageal adenocarcinoma when applied to an independent cohort.

The CNA architecture itself distinguishes normal aging from oncogenic evolution. Healthy lymphocytes and neurons accumulate small, singleton or very-low-frequency events consistent with stochastic damage and imperfect repair, paralleling reported accumulation of private somatic mutations in normal tissues[6,7]. Premalignant and malignant tissues, in contrast, propagate large recurrent CNAs through clonal expansion[37–39]. The aging clock is therefore sensitive to a shared genomic instability axis while preserving this architectural distinction, a property that may prove useful for identifying lesions whose somatic genomes appear disproportionately old relative to patient age. Future lineage-resolved and longitudinal sampling will help disentangle de novo CNA generation, clonal persistence, and cell turnover during hematopoietic aging.

A central finding is that sex-chromosome loss is not a unitary biomarker across sexes. Consistent with prior literature, LOY increased with age in males[14,16,29]; in females, whole-chromosome X loss accumulated even faster than male LOY, with elevated rDNA copy number and accelerated autosomal SNV accumulation in X-loss cells. We observed a transition around ∼30 years, after which SNV burden in both female X-loss cells and male LOY cells exceeded that of non-loss cells from the same donor, consistent with our findings in cancer LOY cells from aged patients[29]. One interpretation is that X-loss occurs in cells experiencing broader genome destabilization, potentially through dosage disruption of X-linked regulators of DNA repair, chromatin maintenance, or nucleolar function (e.g., ATRX, HDAC6, SMC1A)[30–32]. Loss of the inactive X may be partially tolerated while still perturbing higher-order chromatin or nuclear organization, creating a distinct state of genomic stress which is not mirrored by Y loss; that is, X-loss and LOY may differ in fitness consequences. The reduced SNV density observed on the female X chromosome in the Results is also consistent with this framework. Because females carry two copies of chromosome X, one of which undergoes X-chromosome inactivation (XCI) and adopts a condensed heterochromatic state, the heterochromatic inactive X may be less accessible to endogenous mutagenic processes, reducing the per-copy mutation rate[40,41]; additionally, retention of two X copies may impose stronger purifying selection against damaging mutations in essential X-linked genes, preferentially eliminating cells with extensive homozygous variants[42]. These properties may also explain why X-loss, particularly of the inactive copy, carries broader genomic consequences than LOY, as even loss of the largely silenced Xi could disrupt dosage of escape genes or alter three-dimensional nuclear organization. Our prior analysis of this same lymphocyte single-cell dataset supports preferential loss of the inactive X[13], implying that the genomic consequences of female X-loss may arise predominantly from Xi loss rather than random loss of either X chromosome.

Age-associated X loss in females has received substantially less attention than LOY. Our work quantifies X-loss prevalence directly across thousands of lymphocytes spanning infancy to late adulthood and places it in the broader context of the somatic mutational landscape, showing that X-loss cells differ from non-X-loss cells in autosomal SNV burden and rDNA content. By excluding sex chromosomes from clock construction while still recovering sex-specific age trajectories, we further show that female-specific aging structure is embedded in the autosomal CNA background. These observations suggest that X-loss is not merely a passive cytogenetic age marker but identifies a biologically distinct cell subpopulation.

The CNA-based aging clock is best understood as an orthogonal molecular layer complementary to existing clocks. Transcriptomic clocks are sensitive to transient cellular state and remain tissue-specific[20–22]; methylation clocks are among the most accurate chronological-age predictors available but capture regulatory state rather than direct structural genome damage[19]. CNA profiles index cumulative structural genome instability and are likely less susceptible to short-term physiological fluctuation. In addition, our parallel SNV analysis should be interpreted with caution: ultra-CNA was designed for high-throughput copy-number profiling rather than calibrated SNV detection, and low-pass single-cell data are vulnerable to allelic dropout and uneven amplification. Even so, the emergence of SBS1 and SBS5 age associations indicates that the dataset retains meaningful point-mutation signal. Higher-coverage approaches and orthogonal measures (e.g., microsatellite instability, indel burden) will be needed to determine whether combined CNA+SNV models improve age prediction or disease-risk stratification beyond CNA alone.

Several limitations apply. The healthy aging cohort is modest, limiting inference about inter-individual variability and within-person trajectories. Lymphocytes comprise heterogeneous immune subsets, and our analysis did not resolve subset-specific contributions to CNA accumulation or sex-chromosome loss. The female clock was trained on a modest number of donors, and additional cohorts will be needed to establish sex-specific calibration robustness. SNV burden, SBS attribution, and rDNA estimates depend on low-pass single-cell data.

Despite these limitations, our study establishes high-throughput single-cell CNA profiling as a biologically informative framework for studying human genomic aging and demonstrates that sex-stratified calibration is essential for any blood-based molecular aging biomarker. Prospective validation in larger, longitudinal cohorts, integration with immune-subset-resolved profiling, and combination with methylation, transcriptomic, and chromatin-state measurements will clarify how CNA-based age relates to other hallmarks of biological aging and whether blood-based CNA aging signatures can support molecular age monitoring, early detection of age-accelerated disease, and risk stratification for premalignant conditions.

## Materials and Methods

### Single-cell FASTQ read preprocessing (quality control and compression)

Single-cell DNA sequencing data were analyzed from 16 samples (6 female and 10 male) in Tasc-WGS dataset (Liu et al., 2022)[13]. For each cell, a single-end FASTQ file was used as input to the computational pipeline. All analyses were executed on the Texas A&M High Performance Research Computing (HPRC) cluster using SLURM batch scheduling. Raw single-end reads were quality filtered using fastp in multithreaded mode. Cleaned reads were written as gzip-compressed FASTQ files by streaming fastp output directly to pigz for parallel compression, and fastp logs were retained for QC review.

### High-resolution 50-kb binning and normalized coverage estimation

Quality-controlled single-end reads were aligned with Bowtie2 (v2.5.4) using a Bowtie2 index built from the human reference genome GRCh38 (hg38) (Bowtie2-build), with multithreading enabled. Alignments were output in SAM format, and Bowtie2 alignment logs were recorded for each cell. Aligned reads were assigned to genomic bins using a custom Perl script. Briefly, alignments were filtered to remove SAM header lines, reads with an “XS:i” tag, mitochondrial alignments (chrM), and unmapped reads (FLAG 0×4). For remaining reads, genomic positions were mapped to pre-defined dynamic 50-kb bins, and bin counts were accumulated per chromosome. For each bin, a normalized ratio was calculated by scaling bin depth by bin-specific expected counts and by the global read-depth normalization (excluding chromosomes X, Y, and M from the expected total) to generate a per-bin coverage ratio output table. For heatmap visualization, 50-kb bin-level copy-number values were aggregated to 50-kb display bins.

### GC correction, CNV segmentation (CBS), and copy-number estimation

Copy-number analysis was performed in R using the DNAcopy package. Bin-level ratios were first stabilized by adding a pseudocount (+1) and normalizing by the genome-wide mean. GC content for each bin was joined from a reference bin annotation file, and GC bias was corrected using LOWESS regression (span f = 0.05) on log-transformed ratios, producing GC-corrected bin ratios (“lowratio”). Circular Binary Segmentation (CBS) was then applied to log2(lowratio) values with parameters: alpha = 0.01, nperm = 1000, undo.SD = 0.1, and minimum segment width = 5 bins. Segmentation was run on all bins and on a filtered set excluding bins flagged in despike.aisinglecell.spike (bad-bin mask). Segmentation outputs were written to per-cell directories, including raw segmentation files and a per-bin table after bad-bin removal.

Segment means were converted from log2 space back to linear scale (2^seg.mean^). A final per-bin copy-number table was generated by scaling the segmented signal by a constant baseline factor (cn1 = 0.5) and rounding to the nearest integer to yield estimated copy number per bin. For each single cell, copy-number profiles were visualized using the per-bin copy-number output table. From this table, the bin-level copy-number ratio was recomputed as cn.ratio = lowratio/0.5, and the segmented copy-number estimate as cn.seg = seg.mean.LOWESS/0.5; integer copy number was obtained by rounding cn.seg. Per-chromosome figures were generated for chromosomes 1-22, X, and Y by plotting cn.ratio across genomic coordinates within each chromosome and overlaying the segmented track (cn.seg). In addition, a whole-genome profile was generated for each cell by plotting cn.ratio versus absolute genome coordinate and overlaying cn.seg.

### Summary quality reporting and filtering

To summarize QC and processing outputs across cells, MultiQC (v1.27.1) was run on the pipeline output directory to generate an aggregated report. After MultiQC aggregation, single cells were filtered using two quantitative criteria: (i) mapping quality based on the Bowtie2 “overall alignment rate” reported in the MultiQC Bowtie2 summary table; and (ii) signal-noise metrics derived from copy-number profiles. Cells were retained only if they met an alignment-rate threshold (>90%) and passed additional variability-based filters, including a threshold on the median absolute deviation (MAD) of per-bin copy-number ratios and a threshold on the autosomal fraction of bins deviating from copy number 2 (“non-2 ratio”). Cells with missing metric values were excluded prior to filtering. For each sample, lists of retained and excluded cells and a per-cell filtering summary table were generated.

### Copy-number heatmap construction and visualization

For each sample, per-cell copy-number tables were assembled into a matrix with rows corresponding to cells and columns corresponding to genome-ordered 50-kb display bins. Copy-number values were capped at a maximum of 5 for visualization, and cells were hierarchically clustered using single-linkage clustering on pairwise distances to determine the plotting order. Heatmaps were generated with chromosome-aware x-axis labeling, and vertical lines were drawn to indicate chromosome boundaries.

### Single-cell mutation signature analysis by SBS reference set

Single-nucleotide variation mutational signatures were analyzed from donor-level SBS96 catalogs generated from single-cell SNV calls. COSMIC signature assignment was performed using SigProfilerMatrixGenerator and SigProfilerAssignment against the COSMIC SBS reference set[28]. For downstream comparisons across donors, signature activities were normalized by total sequencing reads and overall alignment rate using the same depth-adjusted strategy applied in our SNV summary workflow, thereby reducing technical differences among samples due to sequencing yield and mapping efficiency. Cohort-level analyses included normalized stacked-bar summaries of signature activities, normalized total SBS burden versus age, and signature-specific activity and fractional-contribution plots versus age. Age associations were evaluated using linear regression and Pearson correlation. COSMIC SBS96 classification and signature fitting follow the standard trinucleotide-context framework used for mutational-signature analysis.

### Single-cell SNV calling and SNV burden quantification

For each selected sample, per-cell coordinate-sorted BAM files generated from Bowtie2 alignment were processed individually. SNVs were called from each single-cell BAM using bcftools mpileup followed by bcftools call in multiallelic calling mode. Variant calls were stored in BCF format and then separated into heterozygous and homozygous SNP call sets using genotype-based filtering in bcftools view. Per-cell SNV burden was subsequently quantified from these call sets and normalized by sequencing depth as SNVs per 100,000 reads, with additional adjustment for overall alignment rate to account for variation in sequencing and mapping quality across cells.

### CNA spectrum construction per donor

To summarize recurrent CNAs at the donor level, we generated a donor-specific CNA “spectrum” from the quality-filtered single cells retained for heatmap analysis. For each donor (sample), only autosomal bins (chr1-22) were used. Bin-level integer copy-number calls were collapsed into contiguous CNA segments by merging adjacent 50-kb bins sharing the same copy-number state within each chromosome, and neutral segments (copy number = 2) were removed. CNA segments were then grouped across cells into recurrent events by clustering segments with identical copy-number states on the same chromosome and requiring high reciprocal overlap (≥0.90) between segments; maximal cliques in the resulting overlap graph were used to define event clusters. For each cluster, a consensus interval was defined using the median start and end coordinates across member segments, and cluster prevalence was calculated as the fraction of retained cells carrying the event. Each CNA event was additionally annotated by size (100 kb-1 Mb, 1-10 Mb, 10-40 Mb, >40 Mb), by approximate chromosome-arm positional category (“edge” vs “middle” in each arm), and by mean GC content across the bins spanning the event, which was binned into low/normal/high categories. Donor-level outputs included tables of recurrent CNA clusters and per-cell member events, and summary plots reporting CNA prevalence stratified by event frequency, copy-number category (0, 1, 3, 4, >5), event size, and genomic annotations.

### Bootstrap generation of pseudosamples and CNA clustering

For each donor, we generated 100 bootstrap pseudosamples by resampling a fixed number of single cells (n = 500) with replacement. Because sampling is with replacement, the same original cell can appear multiple times in one bootstrap draw; these duplicate draws were treated as distinct “pseudo-cells” so they contribute independently during downstream clustering. For each bootstrap pseudosample, we reconstructed CNA segments per pseudo-cell (autosomes only) by merging consecutive bins with identical integer copy-number states, excluding copy-neutral segments (CN = 2). Each resulting CNA segment was annotated with segment length and a mean GC content value computed across bins overlapping the segment. Within each bootstrap pseudosample, CNA segments were clustered per chromosome and exact integer copy-number state using a graph-based maximal-clique approach. Two segments were connected (and therefore eligible to fall into the same clique/cluster) only if they satisfied reciprocal overlap ≥90%, i.e., the intersection length divided by each segment’s length was at least 0.90 for both segments. For each resulting cluster, a consensus interval was defined as the median start and end coordinates across member segments, and the cluster frequency was computed as the number of unique (pseudo-)cells contributing at least one segment to that cluster divided by the total number of cells in the bootstrap pseudosample. Each clustered event was then assigned to discrete annotations: (i) GC-content group based on mean GC across the segment (low < 0.37, normal 0.37-0.41, high > 0.41); (ii) copy-number category collapsed to five labels (0, 1, 3, 4, >5) derived from the segment’s integer copy number; (iii) size bin based on segment length (100 kb-1 Mb, 1-10 Mb, 10-40 Mb, >40 Mb); and (iv) a frequency bucket based on the number of cells and cluster frequency (singleton, 2 cells-<1%, 1-5%, 5-20%, 20-50%, >50%). The pipeline produced, per bootstrap pseudosample, a cluster-level table and an event/member-level table; the latter preserves per-cell membership and supports robust annotation of cluster properties when available.

### Feature construction: channelized CNA burden profiles

To train a clock, each bootstrap pseudosample was summarized into a fixed-length feature vector V by aggregating cluster weights over a predefined set of channels formed by the cross-product of GC group (low/normal/high), copy-number category, size bin, and low-frequency buckets (including unique and rare recurrent strata). Each cluster contributed a weight proportional to the number of sampled cells per bootstrap so that totals were comparable across pseudosamples. After channel aggregation, values were normalized within each frequency bucket (i.e., each bucket’s channel values sum to 1 for a pseudosample), which stabilizes bucket-specific composition and reduces confounding by overall event burden differences between pseudosamples. A small pseudocount was added to all entries to ensure numerical stability for divergence-based factorization. Quality control metrics were recorded per pseudosample, including the number of retained clusters and the total retained weight.

### Unsupervised component learning with KL-divergence NMF

We learned latent CNA components using non-negative matrix factorization on the full pseudosample-by-channel matrix V. The factorization was trained across a scan of candidate ranks K (multiple component counts, K = 2, 3, 4, 5, and 6), using a multiplicative-update solver optimized under the Kullback-Leibler divergence loss, which is well-suited to compositional/count-like nonnegative profiles. This yielded a component-by-channel signature matrix (defining each component’s channel pattern) and corresponding pseudosample exposures (how strongly each pseudosample expresses each component). To obtain consistent exposure estimates after learning the component signatures, exposures for each pseudosample were computed by non-negative least-squares projection onto the learned component basis and then converted to exposure fractions (each pseudosample’s exposures sum to 1).

### Donor-level aggregation and selection of the age-associated component

Because each donor contributed many bootstrap pseudosamples, we summarized exposures at the donor level by averaging exposure fractions across all bootstrap pseudosamples from the same donor. Component-age association was evaluated using donor-median exposures rather than individual pseudosamples to reduce within-donor sampling noise and prevent over-weighting donors that contribute many bootstrap replicates.

For each candidate K, we evaluated each component by correlating donor-median exposure with chronological age, computing regression-based goodness-of-fit, and ranking components by an age-association score that favored strong positive association. The best component for that K was defined as the top-ranked age-positive component.

### Model selection across K and clock calibration

Across all scanned K values, we selected the final model based on the performance of the age calibration fitted on donor medians: chronological age was regressed on the donor-median exposure of the selected component using a simple linear model, and the K achieving the best donor-median R² was chosen. The final clock therefore consists of (i) the chosen NMF component signatures, (ii) the identity of the best age-associated component, and (iii) the fitted slope and intercept that map that component’s exposure to age.

### Apparent (in-sample) age prediction for bootstrap pseudosamples

Using the final selected component and linear calibration, we generated apparent predictions for every bootstrap pseudosample used in training. For each bootstrap pseudosample, predicted age was computed as: age = intercept + slope × exposure(best_component), where the exposure is the pseudosample’s exposure fraction for the selected component. Predictions were summarized at the donor level by reporting the median predicted age and empirical variability across bootstrap replicates (e.g., standard deviation and percentile intervals), providing an internal measure of prediction stability driven by cell-resampling variability. Importantly, these predictions are in-sample and therefore reflect apparent performance rather than a fully independent generalization estimate.

### Leave-one-sample-out (LOSO) cross-validation of the CNA aging clock

To evaluate aging clock construction performance, we performed leave-one-sample-out (LOSO) cross-validation in the female cohort. In each fold, one donor was held out entirely (including all of its bootstrap pseudosamples), and the clock was trained only on bootstrap pseudosamples from the remaining donors. For each candidate number of NMF components K ∈ {2, 3, 4, 5, 6}, we fit KL-divergence NMF on the training pseudosamples, selected the age-positive component with the strongest age association using training donor-median exposures, and then calibrated a linear model of age as a function of that component’s exposure using training donor medians. The held-out donor’s pseudosamples were then projected into the learned component space by non-negative least squares to obtain exposure fractions, and ages were predicted using the training-derived linear calibration. Fold-level performance was summarized using predicted ages across all held-out bootstrap pseudosamples for the excluded donor, and repeated across all donors to obtain an internal cross-validated estimate of model performance and stability.

### Application to an external cancer cohort

To apply the trained clock to new samples, each external sample is converted into the same channelized feature vector used in training by reading its cluster data, assigning clusters to the identical channel definition, and performing the same bucket-wise normalization. The external sample vector is then projected onto the learned component signatures using non-negative least squares to obtain exposure fractions, and age is predicted using the stored linear calibration (intercept and slope) applied to the selected component’s exposure.

Overall, this framework yields (i) the learned component signatures (channel weights defining each component), (ii) exposure estimates for each bootstrap pseudosample, (iii) donor-median exposures used for component selection and calibration, and (iv) predicted ages for bootstrap pseudosamples and donor-level summaries. By coupling within-donor bootstrapping with donor-median component selection, the approach explicitly separates biological age association from sampling noise introduced by finite numbers of cells per donor, while KL-NMF provides interpretable, nonnegative CNA patterns that can be inspected at the level of contributing genomic-feature channels.

## Author Contributions

Conceptualization, Y.F. and J.X.; Methodology, Y.F. and B.F.; Validation, Y.F. and B.F.; Investigation, all authors; Visualization, Y.F. and B.F.; Materials, Y.F. and J.X.; Data curation, B.F.; Writing - original draft preparation, Y.F. and B.F.; Writing - review and editing, all authors; Supervision, Y.F. and J.X.; Funding acquisition, Y.F. and J.X.; Project administration, Y.F. and J.X. All authors have read and agreed to the published version of the manuscript.

## Data Availability Statement

Single-cell whole-genome sequencing data for healthy lymphocyte donors were obtained from Tasc-WGS dataset (Liu et al. 2022)[13]. Single-cell sequencing data for the esophageal cancer cohort (normal squamous epithelium, Barrett’s esophagus, and esophageal adenocarcinoma) are deposited in the NCBI Sequence Read Archive (SRA) under BioProject accession PRJNA1439420 and will be made publicly available upon publication of the associated study[29]. Single-cell whole-genome sequencing data for human neocortical neurons were obtained from the NIMH Data Archive (NDA; https://nda.nih.gov/study.html?id=636, Study 2458) as reported by Chronister et al.[11]. All processed data and intermediate files generated during this study are available from the corresponding author upon reasonable request.

## Code Availability

All scripts and computational pipelines developed for this study are available at Zenodo.

## Supplementary Materials

Figures S1 to S13.

