## Supplementary material for "Somatic copy-number alteration signatures reveal sex-divergent aging trajectories accelerated in cancer": Figures S1 to S13.

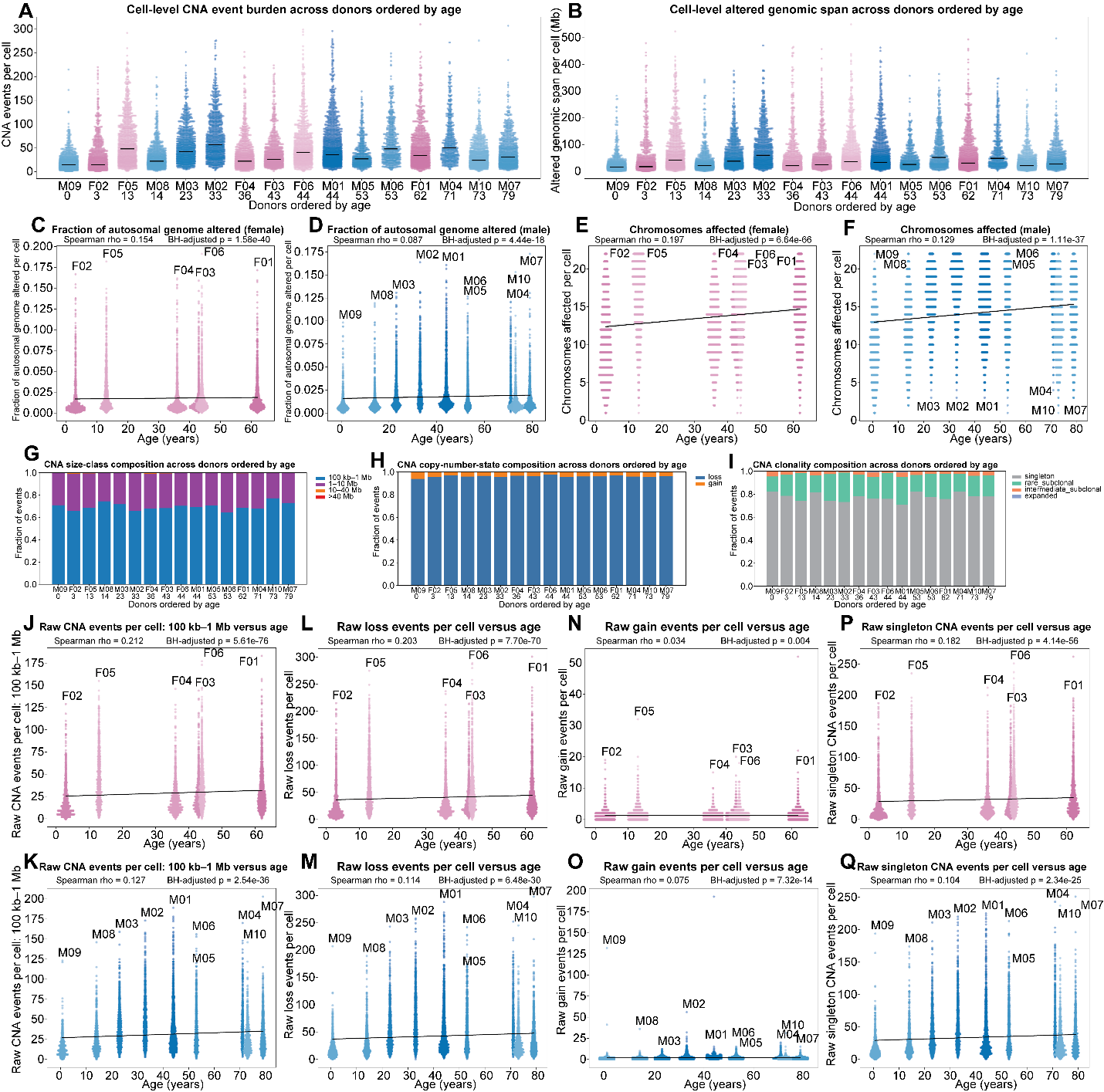

**Figure S1. Single-cell CNA event classification and age associations.**

(A-B) Per-cell CNA event burden (A) and altered genomic span (B) across donors ordered by age. Black bars, donor medians.

(C-D) Fraction of the autosomal genome altered per cell versus age.

(E-F) Number of chromosomes affected by CNAs per cell versus age.

(G-I) Stacked-bar composition across donors (ordered by age) of CNA size class (G), copy-number state (H), and clonality class (I). Fractions normalized within each donor.

(J-K) 100 kb-1 Mb CNA events per cell versus age (J: females, N: males).

(L-M) CNA loss events per cell versus age (K: females, O: males).

(N-O) CNA gain events per cell versus age (L: females, P: males).

(P-Q) Singleton CNA events per cell versus age (M: females, Q: males).

In C-F and J-Q, each point is one cell; pink = female, blue = male; black line, linear fit.

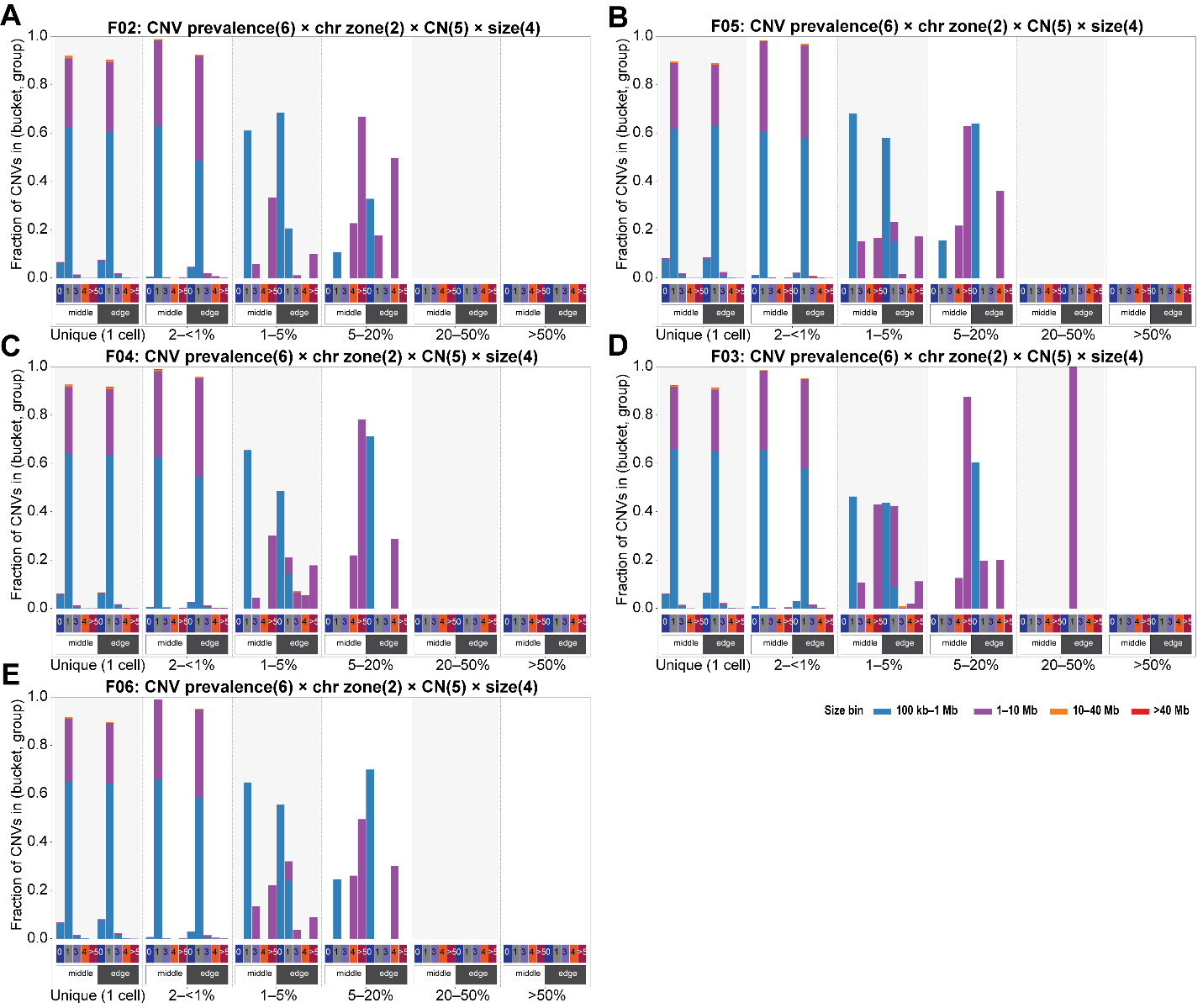

**Figure S2. Arm-zone CNA spectra across the female lymphocyte cohort.**

(A) F02. (B) F05. (C) F04. (D) F03. (E) F06. Layout as in Figure 1H.

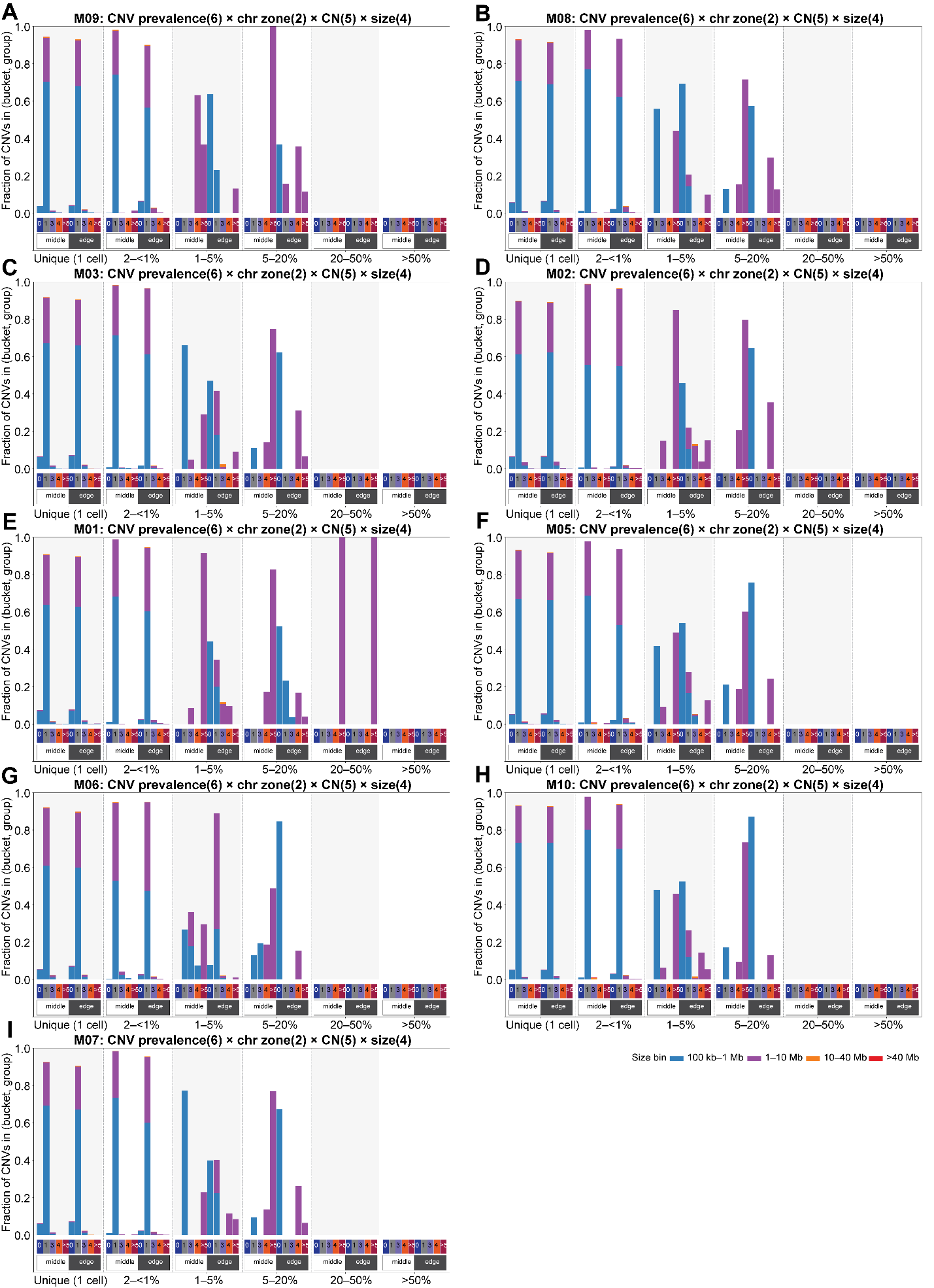

**Figure S3. Arm-zone CNA spectra across the male lymphocyte cohort.**

(A) M09. (B) M08. (C) M03. (D) M02. (E) M01. (F) M05. (G) M06. (H) M10. (I) M07. Layout as in Figure 1J.

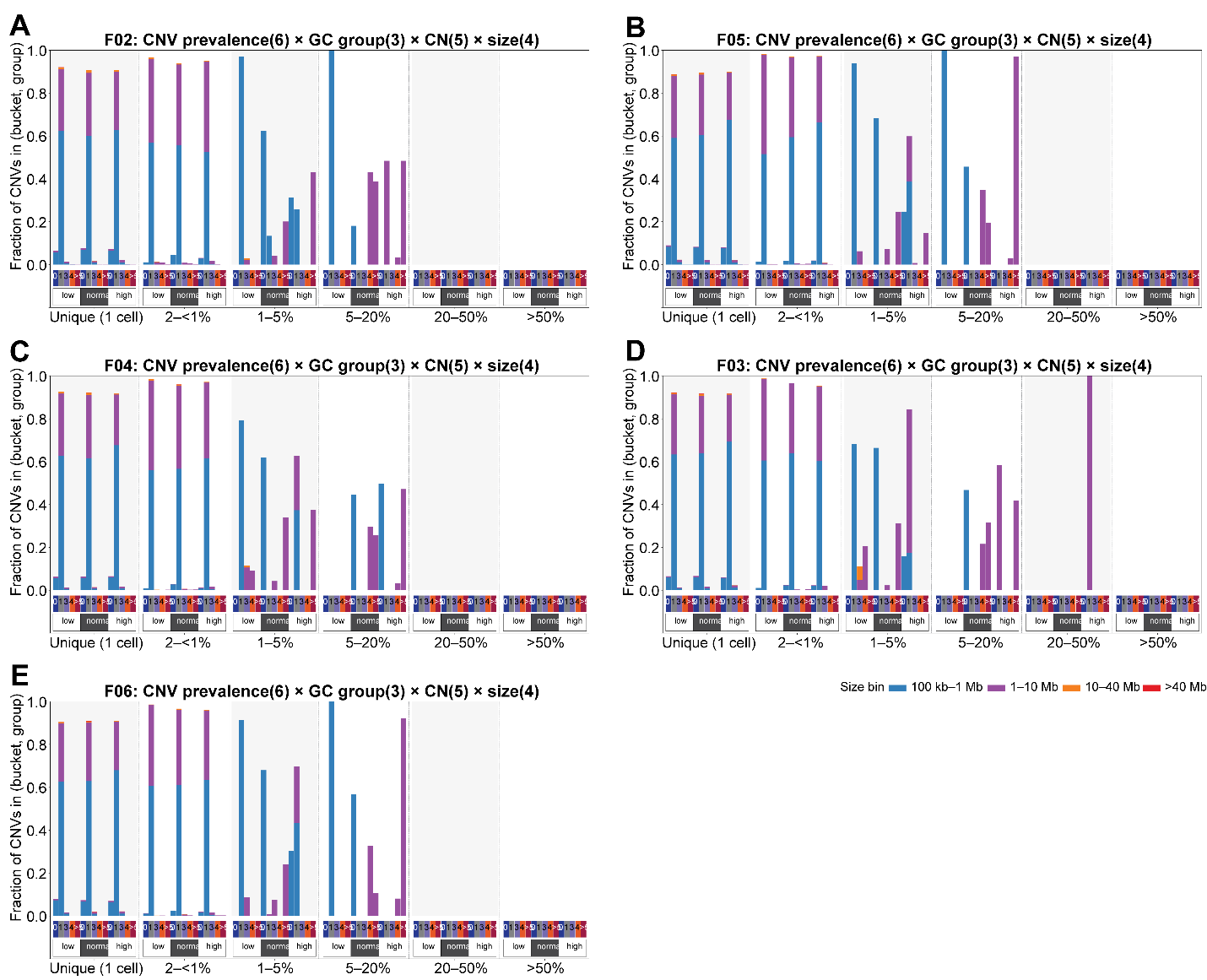

**Figure S4. GC-content CNA spectra across the female lymphocyte cohort.**

(A) F02. (B) F05. (C) F04. (D) F03. (E) F06. Layout as in Figure 1G.

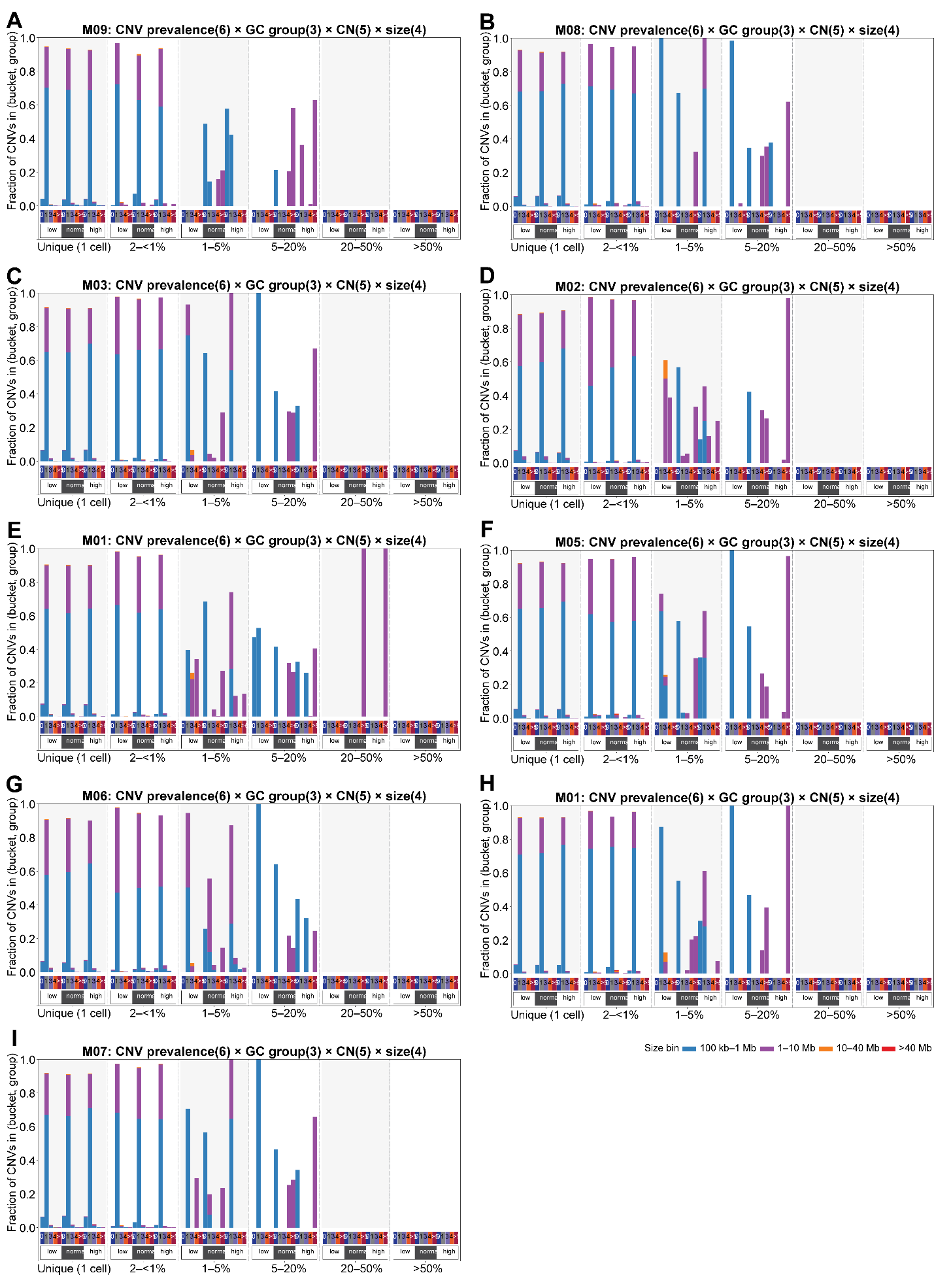

**Figure S5. GC-content CNA spectra across the male lymphocyte cohort.**

(A) M09. (B) M08. (C) M03. (D) M02. (E) M01. (F) M05. (G) M06. (H) M10. (I) M07. Layout as in Figure 1I.

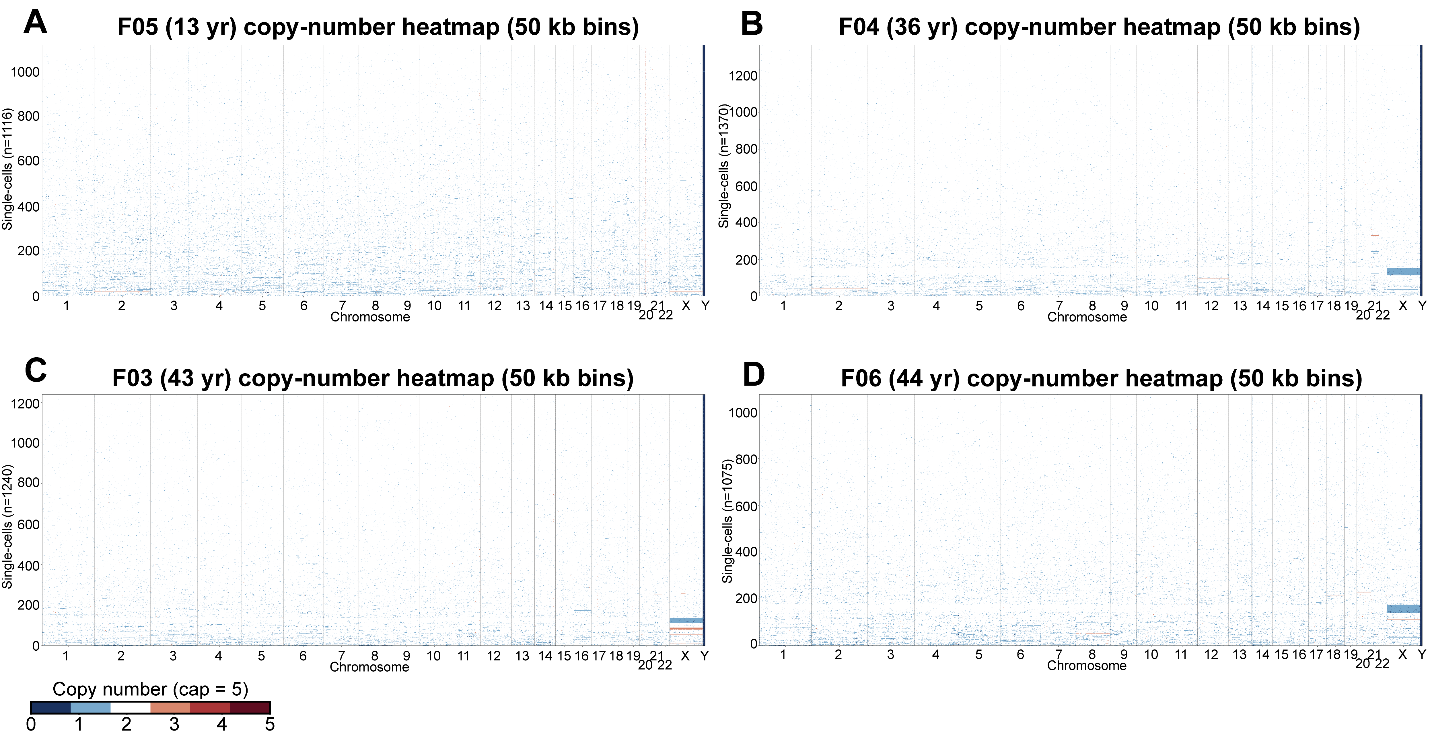

**Figure S6. Single-cell copy-number heatmaps of female donors.**

(A) F05. (B) F04. (C) F03. (D) F06. Layout as in Figure 2A-D.

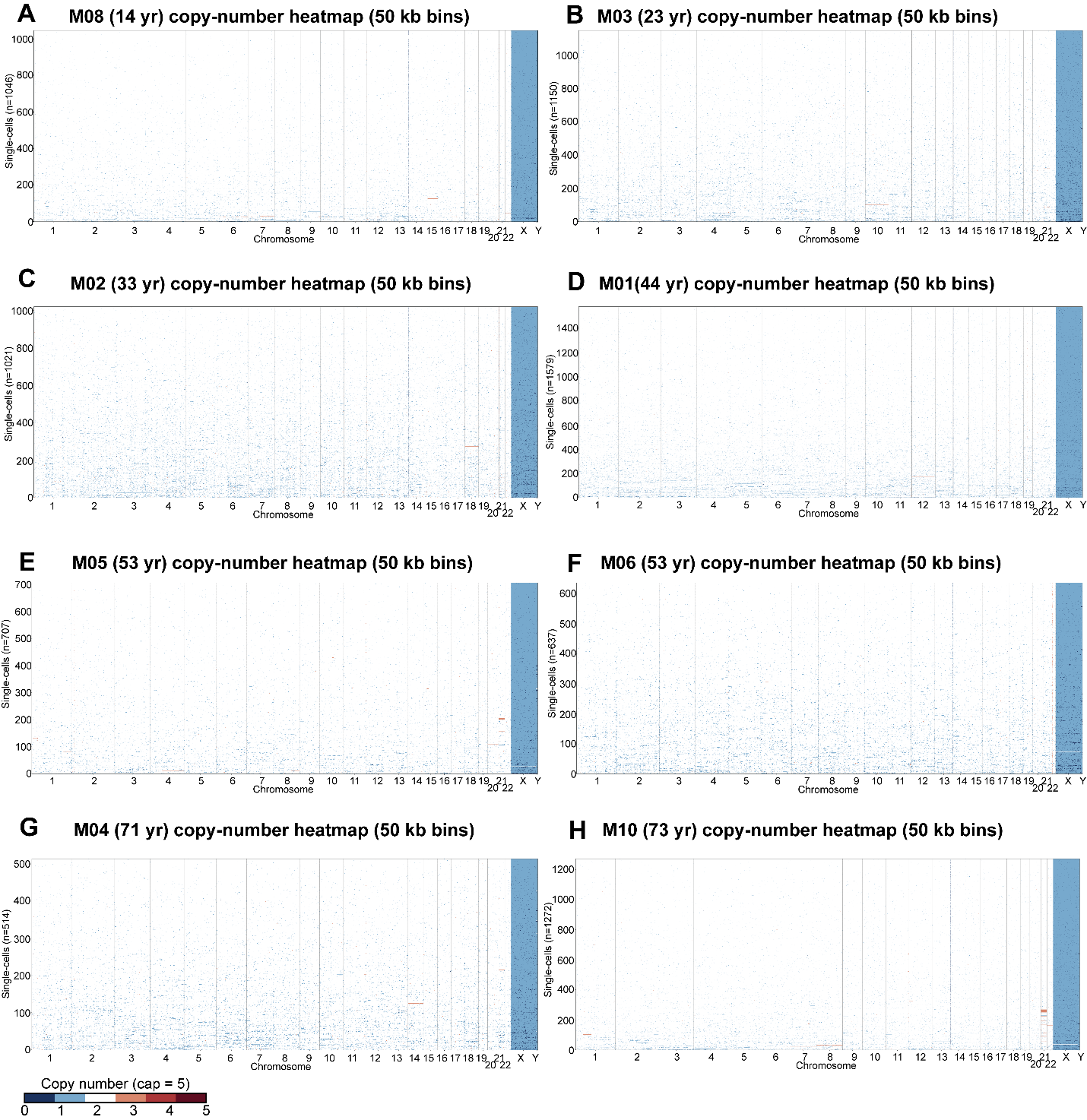

**Figure S7. Single-cell copy-number heatmaps of male donors.**

(A) M08. (B) M03. (C) M02. (D) M01. (E) M05. (F) M06. (G) M04. (H) M10. Layout as in Figure 2A-D.

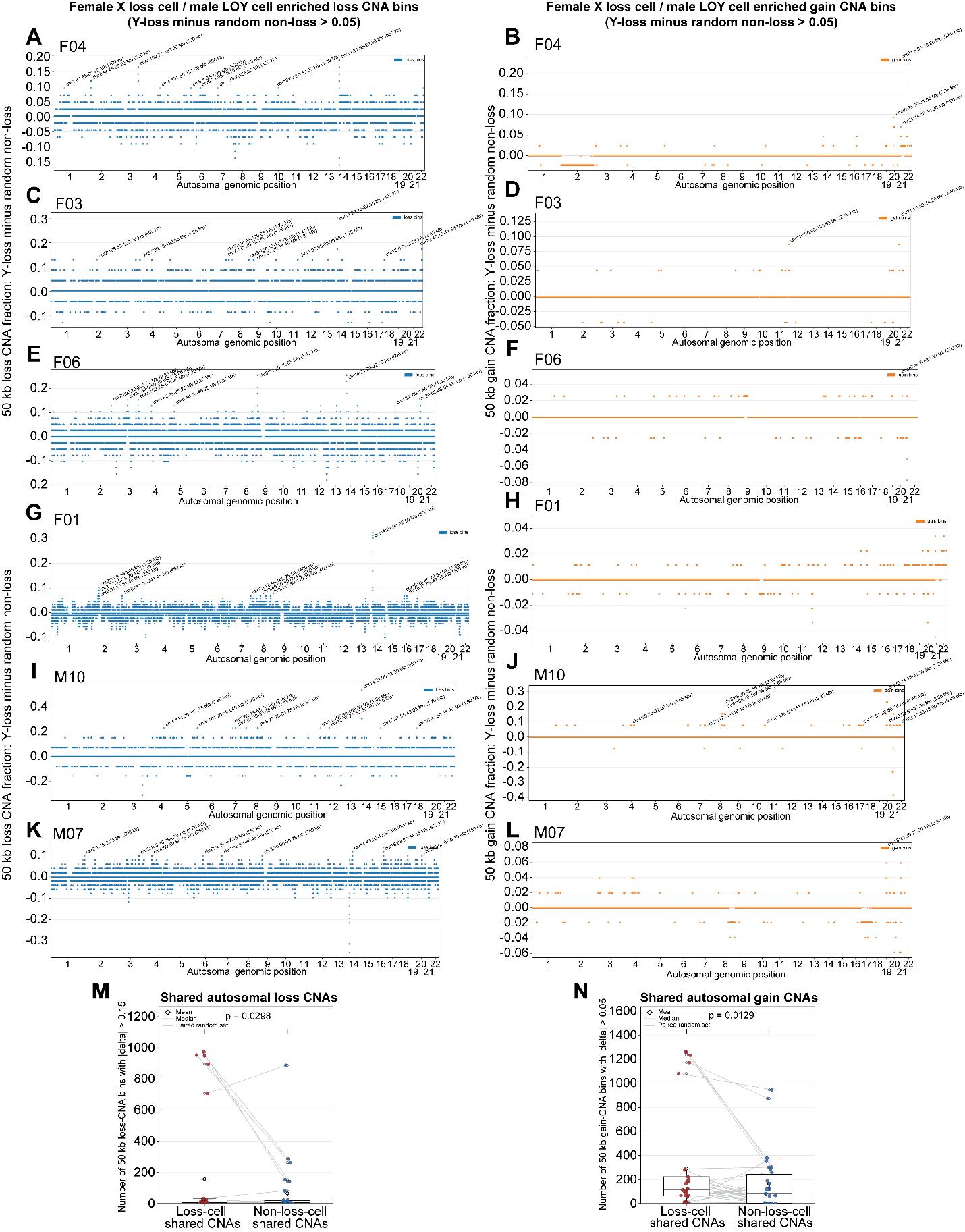

**Figure S8. Recurrent autosomal CNAs in sex-chromosome-loss cells vs. read-depth-matched non-loss cells.**

(A-H) Recurrent loss (A, C, E, G) and gain (B, D, F, H) bins in F04 (A, B), F03 (C, D), F06 (E, F), and F01 (G, H) X-loss cells.

(I-L) Recurrent loss (I, K) and gain (J, L) bins in M10 (I, J) and M07 (K, L) LOY cells.

In A-L, Plots are shown for donors with >10 sex-chromosome-loss cells. Each point is one autosomal 50-kb bin; y-axis, difference in CNA recurrence fraction between loss and read-depth-matched non-loss cells. Blue, loss bins; orange, gain bins. Labeled regions, top recurrent loss- or gain-enriched intervals.

(M-N) Quantification of shared autosomal CNA bins across donor-matched random comparisons. Each point represents one donor/random read-depth-matched comparison. Shared loss-CNA bins were defined using a recurrence-fraction difference cutoff in 0.15, and shared gain-CNA bins using a cutoff of 0.05. (M) Loss-CNA bins. (N) Gain-CNA bins. P values, one-sided paired t-test.

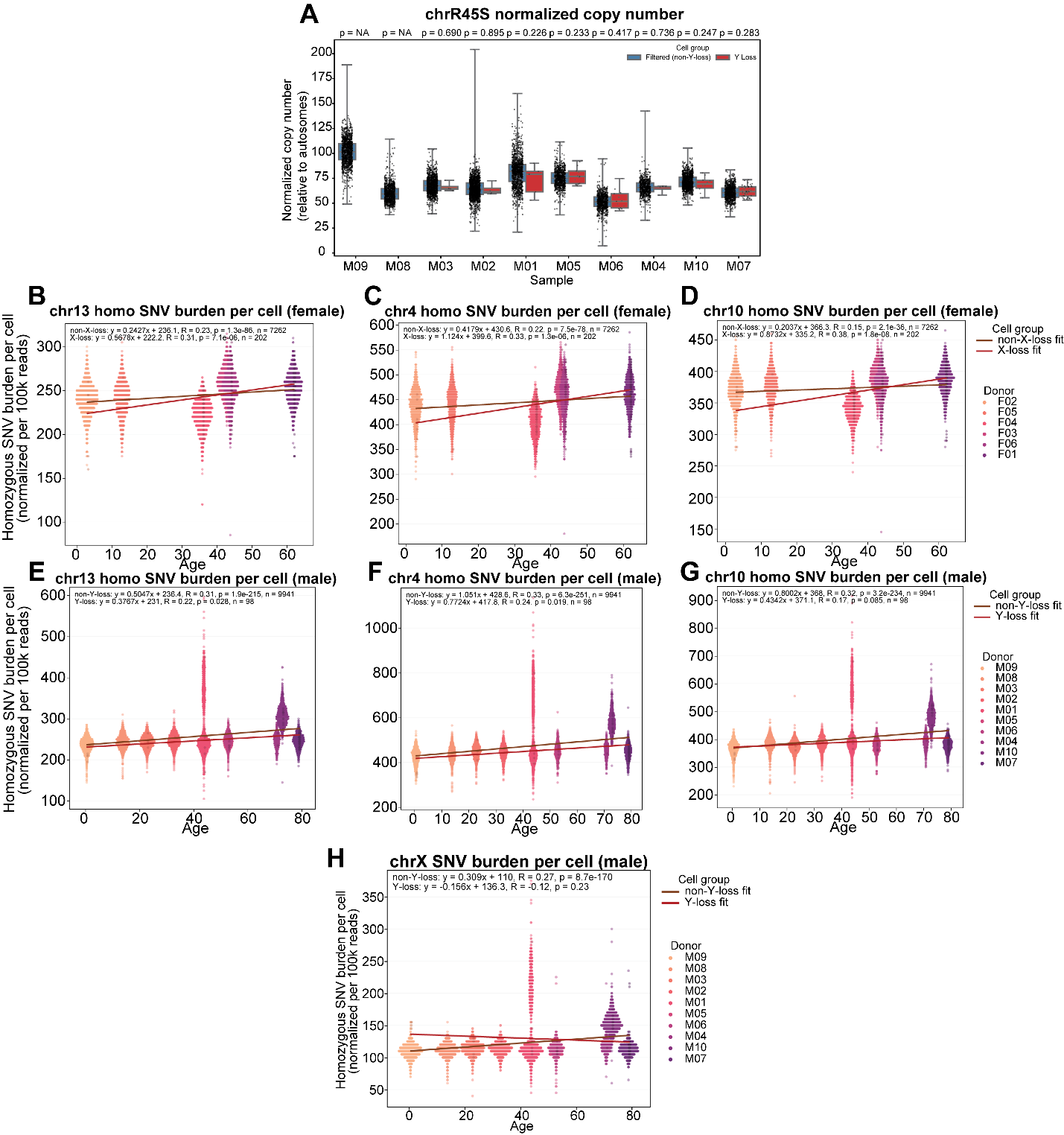

**Figure S9. Chromosomal SNV burden.**

(A) Normalized chrR45S copy number in male LOY versus read-depth-matched non-LOY cells, per donor.

(B-D) Female chr13 (B), chr4 (C), and chr10 (D) homozygous SNV burden per cell versus age.

(E-G) Male chr13 (E), chr4 (F), and chr10 (G) homozygous SNV burden per cell versus age.

(H) Male chrX homozygous SNV burden per cell versus age, stratified by LOY status.

SNV burden normalized per 100,000 reads, adjusted by alignment rate.

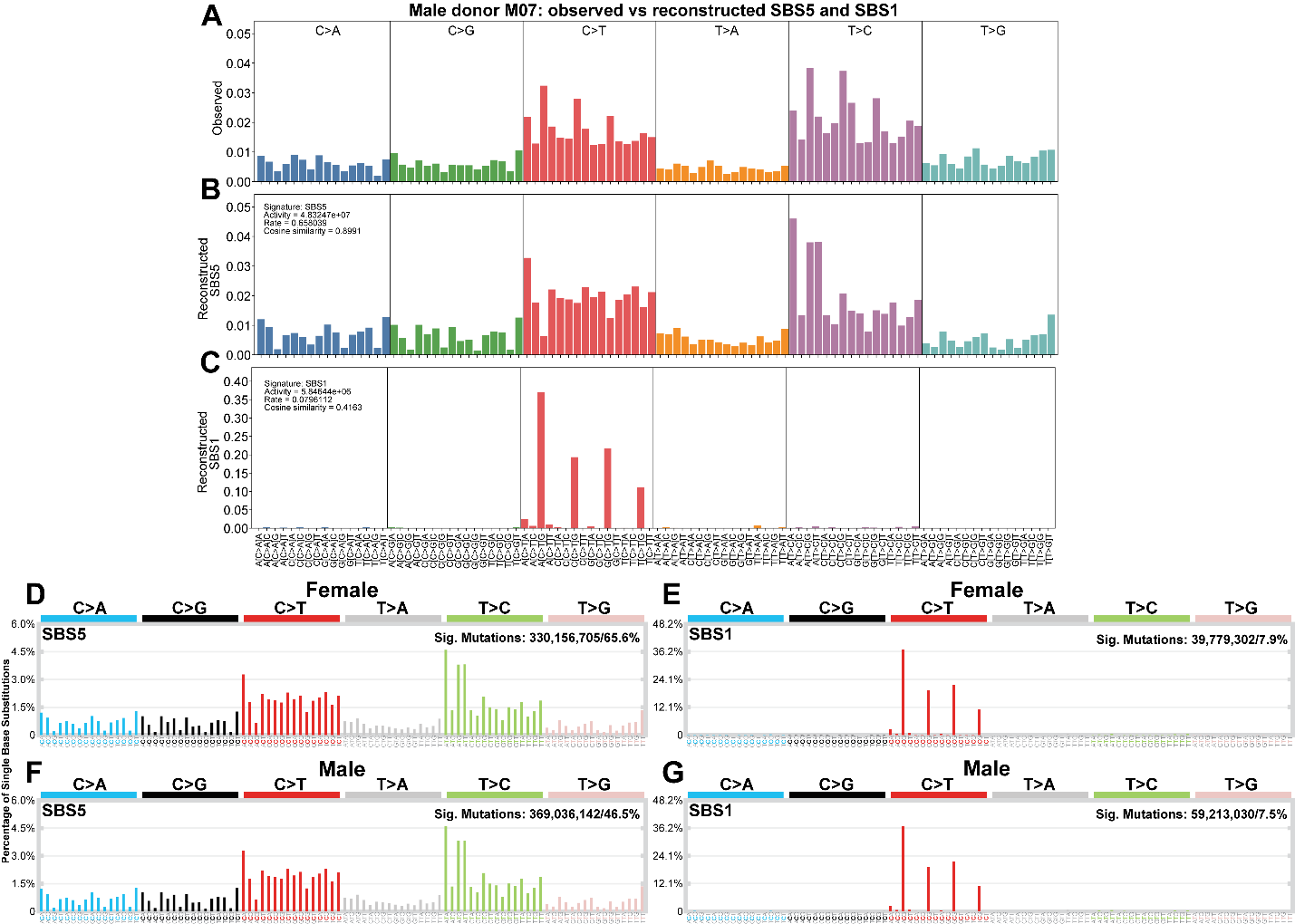

**Figure S10. Donor-level SBS signature decomposition and cohort-wide mutational signature attribution.**

(A) Observed pooled SBS96 spectrum in M07.

(B) Reconstructed SBS96 spectrum from fitted SBS5 contributions in M07.

(C) Reconstructed SBS96 spectrum from fitted SBS1 contributions in M07.

(D-G) Clock-like COSMIC signatures from SigProfilerAssignment. SBS5 (D) and SBS1 (E) in the female cohort; SBS5 (F) and SBS1 (G) in the male cohort. Bars, 96 trinucleotide contexts grouped by substitution class.

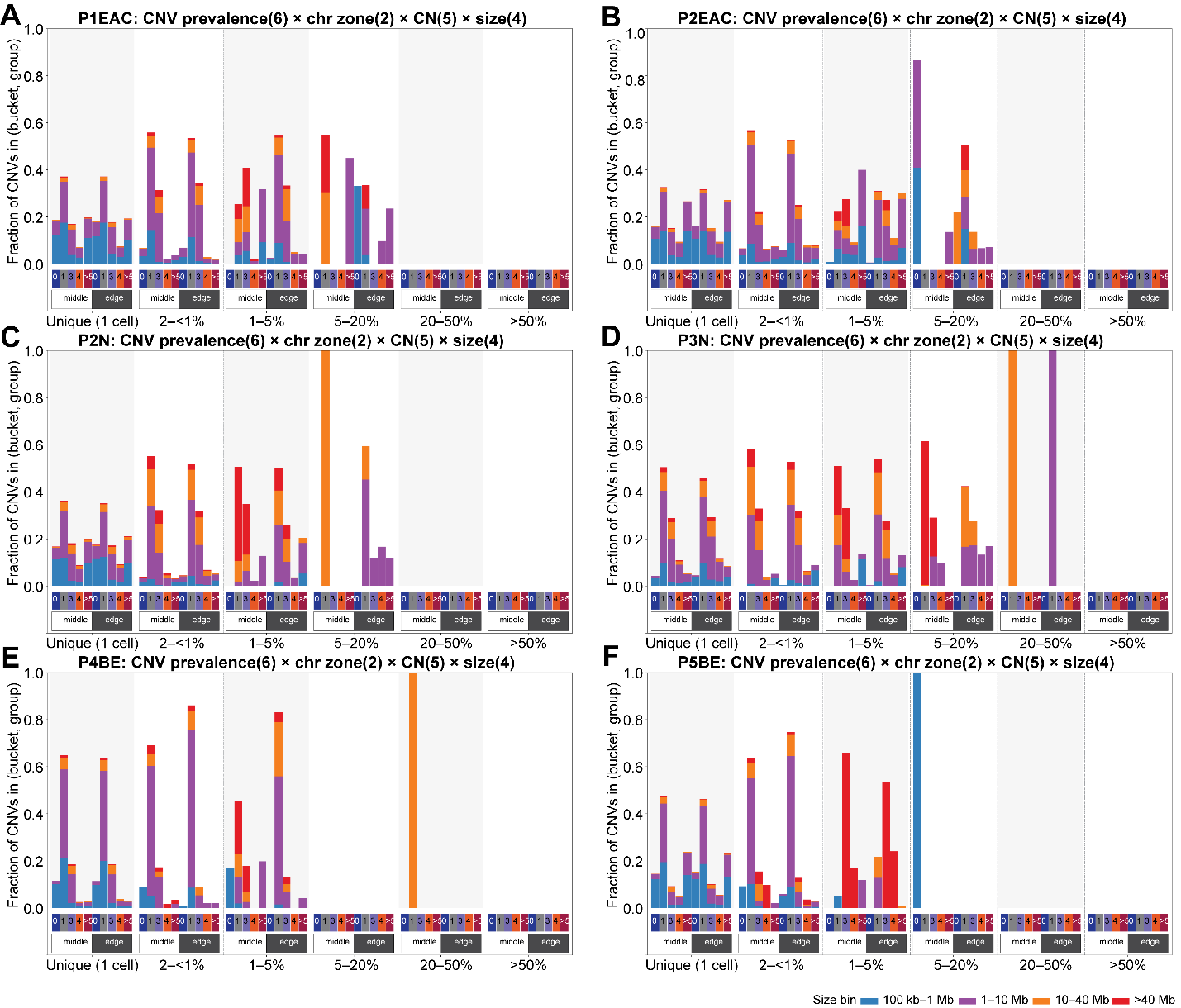

**Figure S11. Arm-zone CNA spectra across the esophageal cohort.**

(A) P1EAC. (B) P2EAC. (C) P2N. (D) P3N. (E) P4BE. (F) P5BE. Layout as in Figure 1H.

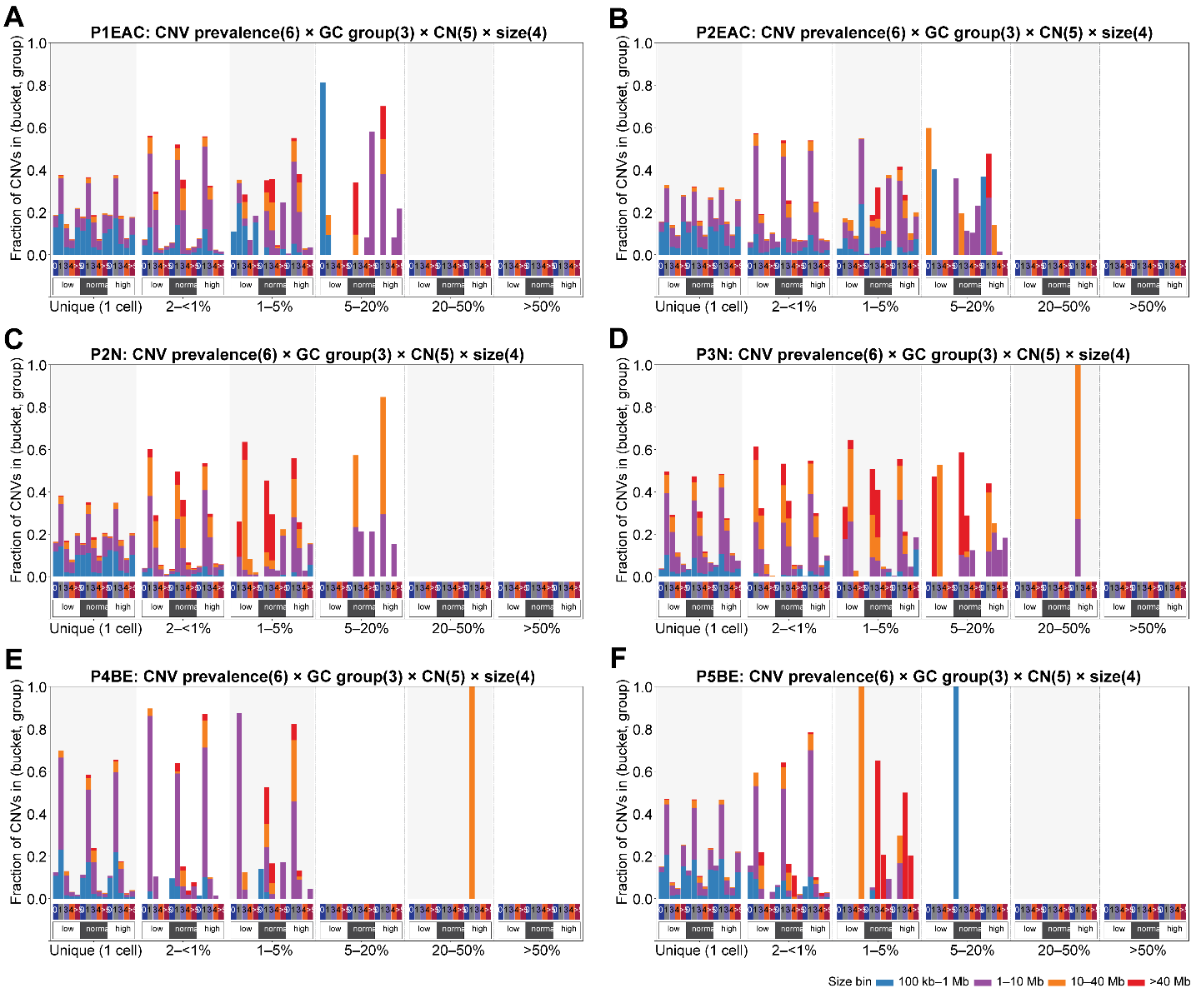

**Figure S12 GC-content CNA spectra across the esophageal cohort.**

(A) P1EAC. (B) P2EAC. (C) P2N. (D) P3N. (E) P4BE. (F) P5BE. Layout as in Figure 1G.

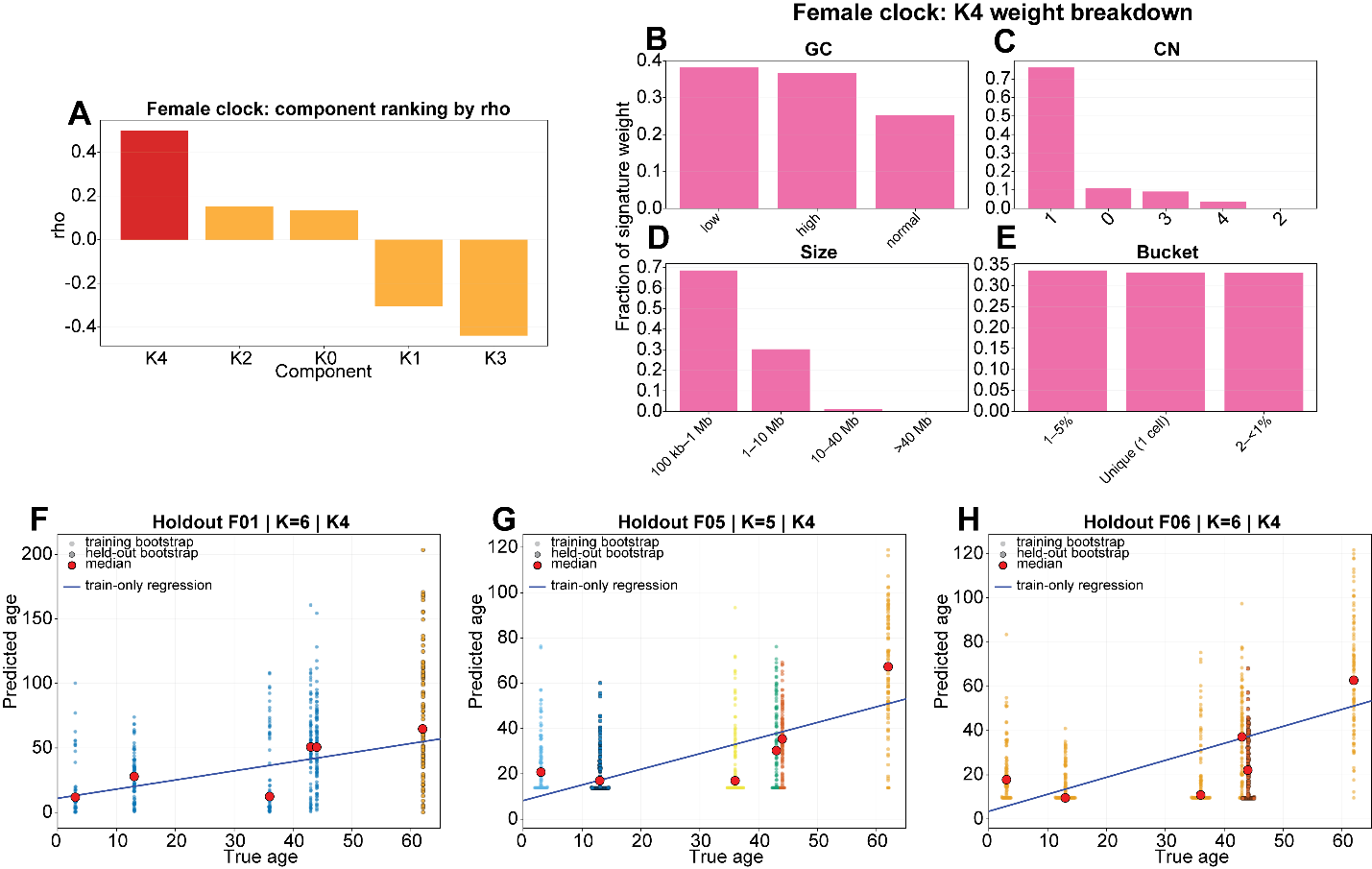

**Figure S13. Component selection and cross-validation of the CNA aging clock.**

(A) Age association of all NMF components in the female training cohort. Bars, Spearman’s rho between donor-level exposure and age; selected aging-clock component highlighted in red.

(B-E) Feature composition of the female aging-clock component: GC content (B), copy-number category (C), CNA size class (D), and clone-frequency bucket (E).

(F-H) Leave-one-sample-out (LOSO) cross-validation in the female cohort. Each point is one bootstrap pseudosample; training bootstraps shown without outline, held-out bootstraps with black outline. Red points, donor medians; blue line, regression on training donor medians.
